# Mint/X11 PDZ domains from non-bilaterian animals recognize and bind Ca_V_2 calcium channel C-termini *in vitro*

**DOI:** 10.1101/2024.02.26.582151

**Authors:** Alicia N. Harracksingh, Anhadvir Singh, Tatiana Mayorova, Brian Bejoy, Jillian Hornbeck, Wassim Elkhatib, Gregor McEdwards, Julia Gauberg, Abdul Taha, Ishrat Maliha Islam, Ted Erclik, Mark A. Currie, Marcus Noyes, Adriano Senatore

## Abstract

PDZ domain mediated interactions with voltage-gated calcium (Ca_V_) channel C-termini play important roles in localizing membrane Ca^2+^ signaling. The first such interaction was described between the scaffolding protein Mint-1 and Ca_V_2.2 in mammals. In this study, we show through various in silico analyses that Mint is an animal-specific gene with a highly divergent N-terminus but a strongly conserved C-terminus comprised of a phosphotyrosine binding domain, two tandem PDZ domains (PDZ-1 and PDZ-2), and a C-terminal auto-inhibitory element that binds and inhibits PDZ-1. In addition to Ca_V_2 channels, most genes that interact with Mint are also deeply conserved including amyloid precursor proteins, presenilins, neurexin, and CASK and Veli which form a tripartite complex with Mint in bilaterians. Through yeast and bacterial 2-hybrid experiments, we show that Mint and Ca_V_2 channels from cnidarians and placozoans interact *in vitro*, and *in situ* hybridization revealed co-expression in dissociated neurons from the cnidarian *Nematostella vectensis*. Unexpectedly, the Mint orthologue from the ctenophore *Hormiphora californiensis* strongly binds the divergent C-terminal ligands of cnidarian and placozoan Ca_V_2 channels, despite neither the ctenophore Mint, nor the placozoan and cnidarian orthologues, binding the ctenophore Ca_V_2 channel C-terminus. Altogether, our analyses suggest that the capacity of Mint to bind Ca_V_2 channels predates pre-bilaterian animals, and that evolutionary changes in Ca_V_2 channel C-terminal sequences resulted in altered binding modalities with Mint.

## Introduction

Ca_V_1 and Ca_V_2 voltage-gated calcium (Ca_V_) channels are high voltage-activated channels that couple membrane excitation with numerous important Ca^2+^-dependent processes including neurotransmitter release in neurons and contraction in muscle [1]. Phylogenetic analyses suggest that these two channel types evolved through gene duplication and diversification very early during animal evolution [2–4], and accordingly, they share common structural features and biophysical properties that distinguish them from the more phylogenetically distant low voltage-activated Ca_V_3 calcium channels [1, 5]. Despite their similarities, Ca_V_1 and Ca_V_2 channels also exhibit some unique and mostly non-overlapping cellular specializations that are broadly conserved in animals. For example, conserved between mammals and invertebrates including arthropods (*i.e.*, the fly *Drosophila melanogaster* and the marine isopod *Idotea baltica*), platyhelminthes (*i.e.*, the flatworm *Dugesia japonica*), nematodes (the worm *Caenorhabditis elegans*), and molluscs (the snail *Lymnaea stagnalis* and the sea hare *Aplysia californica*), Ca_V_1 calcium channels are the primary drivers of cardiac and body muscle contraction [6–13], while Ca_V_2 channels are the primary drivers of neurotransmitter release at fast synapses [14–19]. In part, these differences in cellular function might be supported by evolved differences in instrinsic ion-conducting properties. For example, while both Ca_V_1 and Ca_V_2 channels are subject to negative feedback regulation by the Ca^2+^ sensor protein calmodulin, Ca_V_1 channels exhibit an additional and unique form of regulation by the C-terminal lobe of calmodulin that is insensitive to cytoplasmic Ca^2+^ buffering and accelerates inactivation of the macroscopic current [20–22]. This feature, combined with the generally lower sensitivity of Ca_V_1 channels to voltage-dependent inactivation [23, 24], makes them well suited for roles in cells that exhibit prolonged bouts of depolarization and rises in cytoplasmic [Ca^2+^], as occurs in contracting muscle cells. Mammalian Ca_V_2 channels on the other hand are uniquely subject to negative feedback regulation by G proteins (*i.e.*, Gβγ heterodimers) which become activated by neuromodulatory G protein-coupled receptors [25–28], an important neuromodulatory process that can fine tune synaptic transmission and information flow along neural circuits. Gβγ regulation is also apparent for the Ca_V_2 channel from the mollusc *L. stagnalis* [27–29], but not from the early-diverging animal *Trichoplax adhaerens* [30].

With Ca^2+^ ions generally kept at very low levels in the cytoplasm (*i.e.*, roughly 100 nM) [31], Ca_V_ channel activation results in a discrete and transient domain of cytoplasmic Ca^2+^, with a >1,000-fold increase in concentration from baseline near the channel pore that sharply declines over a distance of 100 nm [32, 33]. A key mechanism for proper activation of Ca^2+^-dependent processes by Ca_V_ channels thus involves their close apposition to calcium-responsive machinery, such that target Ca^2+^-binding sites can become adequately saturated [34, 35]. Interestingly, various protein-protein interactions that distinguish Ca_V_1 and Ca_V_2 channels appear to be broadly conserved, and hence may represent an additional diversification that contributed to the evolution of these two channels types [4, 36, 37]. For example, in mouse and *C. elegans*, Ca_V_1 channels associate with the postsynaptic scaffolding protein Shank, a conserved interaction that permits the calcium channels to activate changes in nuclear gene expression through the transcription factor CREB [38–40]. Here, mammalian Ca_V_1.3 channels and *C. elegans* Ca_V_1 have respective C-terminal short linear motifs (SLiMs) of ITTL_COOH_ and VTTL_COOH_ that are recognized and bound by the PDZ domain of Shank [41], a protein-protein interaction domain named for its shared presence in three proteins: <u>P</u>ostsynaptic density protein 95 kDa (PSD95), *<u>D</u>rosophila* disc large tumor suppressor 1 (Dlg1), and Zonula occludens-1 protein (<u>Z</u>o-1). PDZ domains are commonly found in synaptic scaffolding proteins, where they differentially recognize and bind the extreme C-termini of target proteins via 16 unique ligand specificity classes which are conserved between vertebrates and invertebrates [41].

Importantly, like Ca_V_1 channels, Ca_V_2 channels from vertebrates and invertebrates also exhibit conserved interactions with RIM [42–44] and Mint (a.k.a. X11) [15, 45–49], both via PDZ domains that recognize the extreme C-terminus of Ca_V_2 channels at a conserved consensus ligand sequence of DDWC_COOH_. Indeed, the first report of a PDZ domain binding the C-terminus of any Ca_V_ channel was from Mint [46], a neuronal modular adaptor protein better known for its interaction with amyloid beta precursor protein (APP) [50–53] and its γ-secretase processing enzyme presenilin [54, 55], and hence possible involvement in Alzheimer’s disease [56]. Here, a yeast 2-hybrid assay revealed an interaction between first of two tandem PDZ domains of the rat Mint-1 protein (*i.e.*, PDZ-1) and the extreme C-terminus of the rat Ca_V_2.2 channel bearing a PDZ ligand sequence of DHWC_COOH_ [46]. Subsequently, the *in vitro* interaction between Mint and Ca_V_2 channels from mammals was corroborated though a series of experiments including yeast 2-hybrid, protein pulldown, and nuclear magnetic resonance (NMR) [45, 47, 48]. Furthermore, a direct *in vitro* interaction between Mint and Ca_V_2 was demonstrated for *L. stagnalis* via yeast 2-hybrid [15], and indirectly for chick (*Gallus gallus*) where endogenous Mint protein was pulled down from brain lysates using the Ca_V_2.2 channel C-terminus as bait [49].

Notably, while the physiological function of the RIM-Ca_V_2 channel interaction has been established as important for the proper presynaptic localization of Ca_V_2 channels in mammals [43], *C. elegans* [44], *D. melanogaster* [42], and *A. californica* [16], the physiological significance of the Mint-Ca_V_2 channel interaction remains unclear. In mice, double knockout of the two neuronal Mint genes, Mint-1 and Mint-2, did not disrupt the presynaptic localization of Ca_V_2 channels, despite causing significant disruptions in synaptic morphology and function, and lethality in 80% of newborns [57]. Similarly, genetic ablation of Mint in *C. elegans* caused significant phenotypic defects, including disrupted vulva development and nervous system function, but did not perturb the presynaptic localization of Ca_V_2 channels [18]. Lastly, siRNA downregulation of Mint in *L. stagnalis* did not perturb synaptic transmission in synapses reconstituted *in vitro* mediated by the Ca_V_2 channel [15], altogether suggesting that Mint does not play a role in localizing Ca_V_2 channels to the synapse active zone. Nonetheless, the robust nature of this interaction *in vitro* and its conservation between vertebrates and invertebrates suggests a yet undiscovered neural function.

Spurred by recent observations that many synaptic genes evolved either prior to animals, or are present in early-diverging animals that lack a nervous system and synapses, we set out to explore the phylogenetic origins of Mint and several of its known interacting proteins (*i.e.*, CASK, Veli, Amyloid-β precursor protein, presenilin, and neurexin), and to directly test the *in vitro* interaction between Mint and Ca_V_2 channels from non-bilaterian animals of the phyla Cnidaria (jellyfish, sea anemones, corals), Placozoa (flat aneural animals), and Ctenophora (comb jellies). We demonstrate that the capacity for Mint and Ca_V_2 channels to interact *in vitro* evolved prior to the emergence of bilaterian animals, and that Mint PDZ domains from non-bilaterian animals recognize and bind canonical Ca_V_2 channel C-terminal DDWC-like ligands, as well as the distinct ligand of ETWC_COOH_ present in placozoan Ca_V_2 channels. Furthermore, we provide evidence for this dual binding capacity of Mint which involves distinct structural elements within its C-terminus, comprised of two tandem PDZ domains (PDZ-1 and PDZ-2) and a C-terminal sequence that binds the PDZ-1 ligand-
 binding groove to impose autoinhibition of ligand binding [48] (hereafter referred to as the PDZ-1 Binding Motif or P1BM).

## Results

### Phylogenetic analyses point to deep conservation of the Mint PDZ-1/Ca_V_2 calcium channel interaction

Ca_V_2 channels are unique to animals, having emerged alongside Ca_V_1 channels via an ancient gene duplication event [2, 4]. A protein phylogeny of select Ca_V_2 channel sequences produced a branching pattern mostly consistent with the expected species phylogeny rooted on the most divergent orthologues from the ctenophore species *Mnemiopsis leidyi* and *Hormiphora californiensis* (Figure 1A). The three vertebrate Ca_V_2 channel isotypes, P/Q-type (Ca_V_2.1), N-type (Ca_V_2.2), and R-type (Ca_V_2.3) form a strongly supported monophyletic clade reflecting their proposed emergence through more recent gene/genome duplication events that occurred within the phylum Chordata, after tunicates (*e.g.*, *Ciona intestinalis* and *Styela clava*) and cephalochordates (*Branchiostoma belcheri*) diverged. Consistent with previous reports, the well-supported monophyletic clades of Ca_V_2 channels from cnidarians and platyhelminthes reflect independent duplication events that likely gave rise to Ca_V_2a, Ca_V_2b, and Ca_V_2c in cnidarians, and Ca_V_2A and Ca_V_2B in platyhelminthes [2, 4, 36].

**Figure 1.**
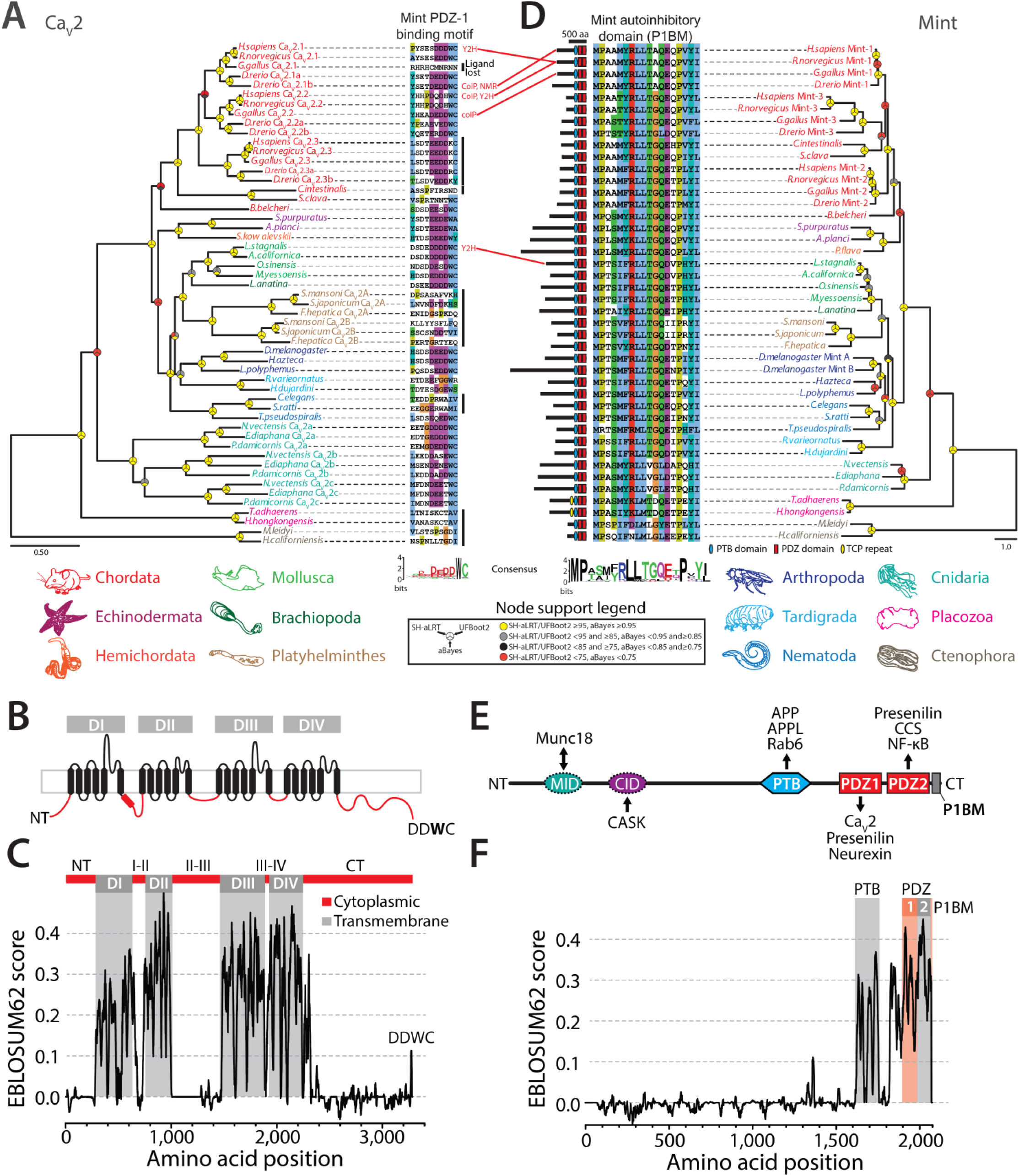
The structural elements required for the Mint-Ca_V_2 interaction are conserved in cnidarians and bilaterians. **(A)** Maximum likelihood protein phylogeny of Ca_V_2 channels from selected animals, with names color-coded according to phylum. Sequences of the extreme C-termini of the channels are shown on the right of the phylogenetic tree, revealing a deeply conserved PDZ ligand motif with a consensus sequence of DEDDWC. Species for which a direct interaction between Ca_V_2 channels and Mint have been experimentally determined are indicated in red text to the right of select Ca_V_2 channel C-terminal sequences (coIP: co-immunoprecipitation; Y2H: Yeast 2-Hybrid; NMR: Nuclear Magnetic Resonance spectroscopy). **(B)** Illustration of the Ca_V_2 channel transmembrane topology, depicting four homologous repeat domains (DI-DIV) each with 6 transmembrane alpha helices (in black), as well as the intracellular N- and C-termini and cytoplasmic linkers that separate the four domains (in red). **(C)** EMBOSS Plotcon alignment similarity plot for single Ca_V_2 channel representatives selected from each phylum from the tree depicted in panel *A*, revealing extensive divergence in amino acid sequences within the cytoplasmic regions (linkers and N- and C-termini), but strong conservation within transmembrane regions of Domains I-IV. Notably, the conserved DDWC ligand sequence appears as a short linear motif at the extreme end of the highly divergent and structurally disordered C-terminus. **(D)** Maximum likelihood phylogeny of Mint proteins from select animals. The lengths of the various Mint proteins are indicated with black bars on the left of the phylogenetic tree. The symbols on the black bars denote ubiquitously conserved PTB and tandem PDZ domains (i.e., PDZ-1 and PDZ-2), as well as unique tetratricopeptide (TCP) repeat domains in the placozoan orthologues [77], as predicted with InterProScan. Also to the left of the tree is an alignment of the extreme C-terminal sequences of the various Mint proteins, revealing deep conservation of the auto-inhibitory domain (P1BM) that associates with the ligand binding pocket of PDZ-1 causing ligand binding auto-inhibition [48]. For panels A and D, node support values generated with SH-aLRT, UFBoot2, and aBayes are indicated by symbols according to the provided legend. **(E)** Illustration of the various domains found within the Mint protein that are involved in interactions with other proteins (MID: Munc-18 Interacting Domain; CID: CASK Interacting Domain; PTB: PhosphoTyrosine-binding Domain; PDZ domains 1 and 2: PDZ1 and PDZ2; P1BM: PDZ-1 auto-inhibitory peptide). The black arrows indicate domain-ligand relationships, with the head pointing towards the ligand. **(F)** EMBOSS Plotcon alignment similarity plot for single Mint representatives selected from each phylum from the tree depicted in panel B, revealing a highly divergent N-terminus, but strong sequence conservation within the C-terminal PTB domain, the tandem PDZ domains, and the P1BM auto-inhibitory domain.

Alignment of the extreme C-terminal protein sequences of Ca_V_2 channels reveals a conserved amino acid motif with a consensus sequence of DDWC_COOH_, shared between cnidarian and bilaterian homologues (Figure 1A). This PDZ ligand motif can be classified as a SLiM, preceded by a long stretch of highly divergent amino acids that contribute to a mostly disordered Ca_V_2 channel C-terminus (Figure 1B and C). The most invariable amino acid within this motif is the tryptophan residue (W) in the second last position, a common feature of PDZ ligands being present in 5 of the 16 defined specificity classes [41]. Nonetheless, the presence of a distal C-terminal cysteine residue indicates that this motif is a non-canonical PDZ domain ligand. Apparent is the independent loss of DDWC-like motifs in several lineages of Ca_V_2 channels. Within chordates, the chick (*Gallus gallus*) Ca_V_2.1 channel and the *C. intestinalis* Ca_V_2 channel completely lost this motif, the former attributed to the loss of a C-terminal exon that encodes the distal portion of the Ca_V_2 channel C-terminus in chick and perhaps all birds [4, 58]. Notably, vertebrate Ca_V_2.3 channels bear cationic lysine/arginine (K/R) residues rather than tryptophan in the second last position, which likely disrupts PDZ domain binding since rat Mint-1 does not bind the human Ca_V_2.3 C-terminus *in vitro* [46], and mutation of the *L. stagnalis* Ca_V_2 channel DDWC motif to DDKC disrupts its interaction with Mint [15]. Given the conservation of DDWC-like C-terminal SLiMs in cnidarian and bilaterian Ca_V_2 channels, it appears that platyhelminthes and the nematode species *Caenorhabditis elegan*s and *Strongyloides ratti* independently lost the DDWC_COOH_ motif. However, since the Ca_V_2 channel from the early-diverging nematode *Trichinella pseudospiralis* does bear a DDWC-like SLiM (EDWC), loss of this motif only occurred in a subset of nematode species. Lastly, we note a complete absence of DDWC-like motifs in Ca_V_2 channel C-termini from placozoans and ctenophores, the former lacking a nervous system and synapses [59], and the latter comprising the most divergent animal phylum to possess Ca_V_2 channels, and which interestingly, are proposed to have independently evolved a nervous system and synapses [60].

### Phylogeny of Mint and several key interacting proteins

To better understand the phylogenetic properties of Mint, we mined select genomes and transcriptomes to generate a maximum likelihood phylogeny which was rooted by the ctenophore orthologues (Figure 1D). Given the absence of Ca_V_2 channels in poriferans (sponges) [2, 4], we excluded them from our analysis despite identifying bona fide Mint homologues for several species including *Amphimedon queenslandica*. Like Ca_V_2 channels, most invertebrates included in our analysis seem to possess only single Mint orthologues, with independent duplications apparent for vertebrates (*i.e.*, Mint 1 to 3) and *D. melanogaster* (Mint A and B). Analysis of the identified Mint protein sequences with InterProScan [61] reveals complete conservation of a predicted <u>P</u>hosphotyrosine-<u>B</u>inding (PTB) domain, followed by two tandem PDZ domains (PDZ-1 and PDZ-2) near the C-terminal end of the protein (Figure 1D and E). Alignment of the C-termini of the analyzed Mint sequences reveals strong sequence conservation (Figure 1D), notable because this sequence serves an auto-inhibitory function on the Mint protein by binding to the PDZ-1 ligand recognition groove and thus inhibiting ligand binding [48]. In contrast to the conserved C-terminus, the N-terminus of Mint is quite variable in length (Figure 1D) and amino acid sequence, evident in a plot of global sequence conservation among single representative Mint orthologues from each phylum included in our analysis (Figure 1F). This strong sequence divergence is notable because the N-terminus of Mint bears the CASK Interaction Domain (CID), conserved between vertebrates (Mint-1 only), *D. melanogaster*, and *C. elegans* [62], which contributes to the formation of a tripartite complex with the proteins CASK and Veli [63–65]. The N-terminus of Mint also harbours the Munc18 Interaction Domain (MID), a protein-protein interaction that associates Mint function with vesicle exocytosis and hormone secretion [66]. Altogether, the fact that our alignments failed to identify conserved CID and MID signatures suggests these elements are either highly divergent in protein sequence, or are absent in early-diverging, non-bilaterian Mint orthologues. Nonetheless, we note conservation of both CASK and Veli in all metazoans included in our study (Figure 2A and B), most bearing canonical predicted domain architectures including a Calmodulin Kinase (CaMK) domain at the N-terminus of CASK shown in crystal structures to associate with the Mint CID motif (Protein Data Bank [PDB] accession number 6LNM), as well as Lin-2/Lin-7 (L27) domains, important for the interaction between CASK and Veli [64].

**Figure 2.**
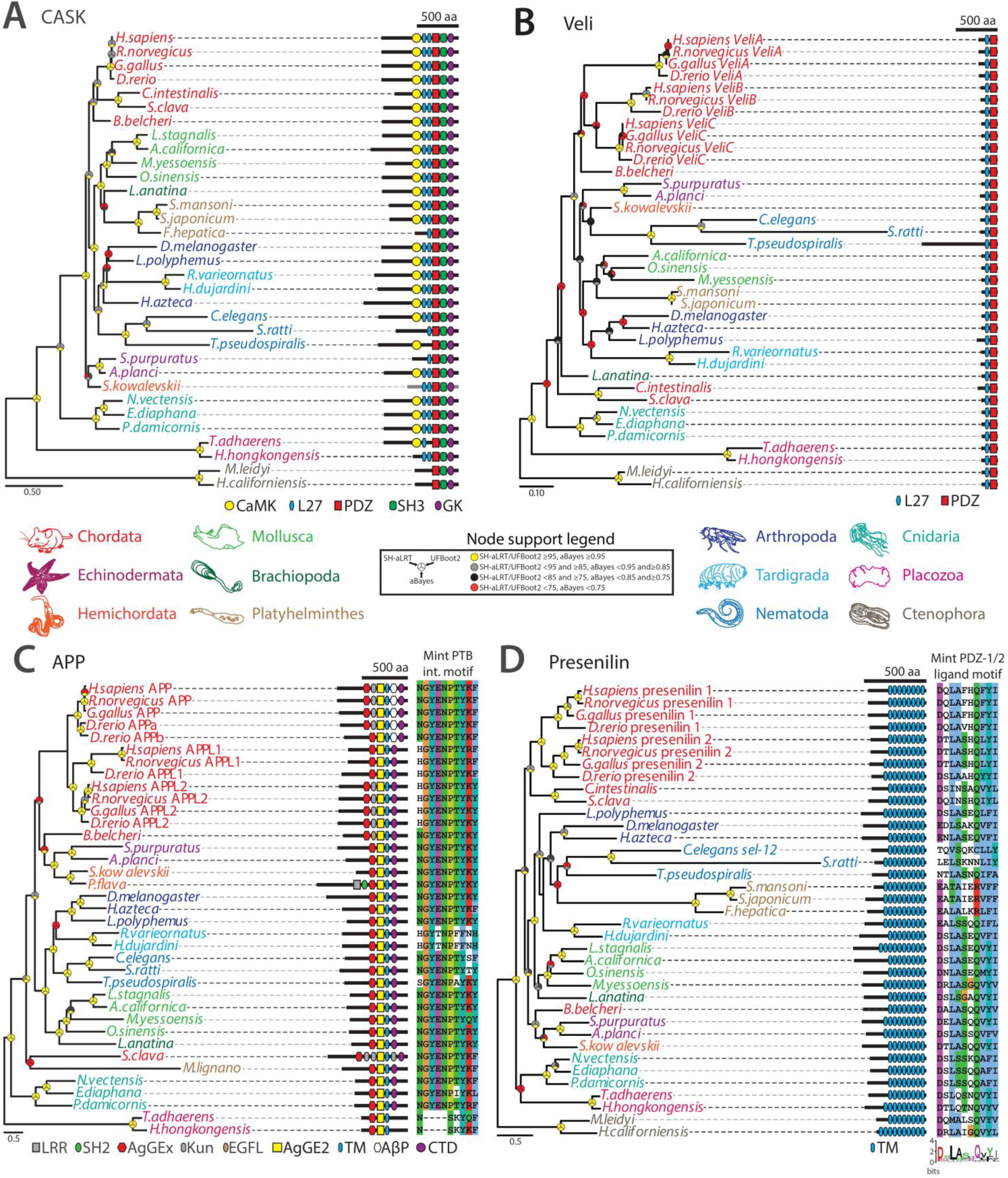
Phylogenetic properties of several key proteins that interact with Mint. **(A)** Maximum likelihood protein phylogeny of CASK homologues from various animals. Protein lengths are indicated with black bars, and domains predicted with InterProScan are illustrated (CaMK: Ca^2+^/calmodulin-dependent protein kinase domain; L27: Lin-2/Lin-7 domain; PDZ: PSD95/Dlg1/zo-1 domain; SH3: Src Homology 3 domain; GK: guanylate kinase domain). **(B)** Maximum likelihood protein phylogeny of Veli homologues from various animals. Protein lengths are indicated with black bars, and domains predicted with InterProScan are illustrated (L27: Lin-2/Lin-7 domain; PDZ: PSD95/Dlg1/zo-1 domain). **(C)** Maximum likelihood protein phylogeny of amyloid β precursor proteins (APP) and amyloid β-like (APPL) homologues from various animals, colour-coded according to phyla. C-terminal sequences are shown on the right, illustrating deep conservation of the YENPTY motif which is bound to by the Mint PTB domain. No ctenophore homologues were identified, and the placozoan and tardigrade sequences lack the canonical YENPTY motif. The lengths of the APP and APPL proteins are indicated with black bars, and protein domains predicted with InterProScan are illustrated (LRR: leucine-rich repeat domain; SH2: Src Homology 2 domain; AgGEx: amyloidogenic glycoprotein, extracellular domain; Kun: pancreatic trypsin inhibitor Kunitz domain; EGFL: EGF-like domain; AgGE2: amyloidogenic glycoprotein, E2 domain; TM: transmembrane helix; AβP: amyloidogenic glycoprotein, amyloid-beta peptide; CTD: beta-amyloid precursor protein C-terminal). **(D)** Maximum likelihood protein phylogeny of presenilin homologues from various animals, colour-coded according to phyla (legend below). An alignment of extreme C-terminal protein sequences is shown on the right, illustrating deep conservation of the ligand sequence for the Mint PDZ-1 domain. The lengths of the presenilin proteins are indicated with black bars, and transmembrane helices (TMs) predicted with Phobius are illustrated. For all panels, node support values generated with SH-aLRT, UFBoot2, and aBayes are indicated by symbols according to the provided legend.

The deep conservation of the Mint PTB, tandem PDZ, and C-terminal auto-inhibitory PDZ-1 Binding Motif (P1BM), suggests that the various interactions mediated by these domains might also be conserved. The PTB domain, for example, mediates protein-protein interactions with several partners including Amyloid Precursor Protein (APP) [50–53], Amyloid Precursor Protein-Like (APPL) [67], and the Golgi transport protein Rab6 [68] (Figure 1E). We therefore sought to survey the phylogeny of some of these proteins, with a primary focus on exploring conserved motifs associated with Mint interactions, and predicted domain architectures. Interestingly, our analysis revealed that APP and APPL homologues are broadly conserved in animals (Figure 2C), finding homologues in all examined species with the exception of ctenophores, most bearing a YENPTY motif that forms a beta strand which integrates into a binding pocket of the Mint PTB domain [50, 69]. We only found single APP homologues in invertebrates, but a strongly supported monophyletic clade of vertebrate homologues consistent with gene duplications giving rise to APP, AAPL1, and APPL2 (Figure 2C). As noted, the first PDZ domain of Mint (PDZ-1), recognizes and binds the Ca_V_2 channel DDWC motif [15, 46, 47, 70]. Mint PDZ-1, as well as PDZ-2, also recognize the C-terminus of the transmembrane protein presenilin [48, 54, 55] which is part of the γ-secretase complex that regulates target proteins including APP via proteolytic processing [71], and PDZ-1 binds the trans-synaptic cell adhesion protein neurexin [45, 47]. Phylogenetically, all examined species possess presenilin homologues with vertebrates uniquely duplicating the gene to give rise to presenilins 1 and 2 (Figure 2D).

Additionally, all of the identified presenilin homologues possess 8 to 10 predicted transmembrane helices, as well as a conserved hydrophobic C-terminus with a consensus sequence of DSLASHQVYI_COOH_ (Figure 2D), reflecting deep conservation of a putative Mint PDZ-1/PDZ-2 ligand sequence as defined for mammalian Mint 1 and 2 *in vitro* [48, 54, 55]. We also conducted a phylogenetic analysis of synaptic neurexin homologues (referred to as neurexins 1 to 3 in vertebrates and neurexin-1 in invertebrates, collectively referred to as neurexin-1 genes in this study), which belong to a larger group of cell adhesion proteins collectively referred to as the neurexin family [72] (Figure 3). For our analysis, we focused on homologues bearing domain signatures of two LamG2 domains flanked by EGF-like domains, which are unique to neurexin-1. Also, we only included one of each of the three types of neurexin homologues previously identified in *Nematostella vectensis* (*i.e.*, neurexins alpha 1, delta 1, and epsilon 1), recently shown to possess at least 10 neurexin homologues [73]. Like the other analyzed genes, the phylogenetic analysis suggests that neurexin-1 duplicated in vertebrates to produce neurexins 1 to 3, and separately in spiralian species within the phyla Mollusca, Brachiopoda, and Platyhelminthes, the chelicerate arthropod *Limulus polyphemus*, and both placozoans and cnidarians. In molluscs and brachiopods, one of the duplicated neurexin-1 paralogues retained a canonical predicted domain architecture comprised of repeating EGF-LamG2-LamG2 domains and a C-terminal transmembrane helix, while the other incorporated a unique C-terminal WD40 domain and an extended arrangement of LamG2 domains near the N-terminus. This derived isotype from the brachiopod species *Lingula anatina* also possesses a unique set of 9 predicted tandem fibronectin type 3 domains that are associated with cell adhesion and migration. A similar phenomenon is apparent for the duplicated neurexin-1 paralogues from platyhelminths, with one isotype retaining a more canonical domain arrangement, and the other being considerably shorter in length with less predicted LamG2 and EGF domains. With respect to putative C-terminal PDZ ligands, it is notable that most of the examined neurexin homologues exhibit conserved C-terminal KE[W/Y]YV_COOH_ motifs (Figure 3), shown to associate with the PDZ-1 domain of rat Mint-1 via surface plasmon resonance [47].

**Figure 3.**
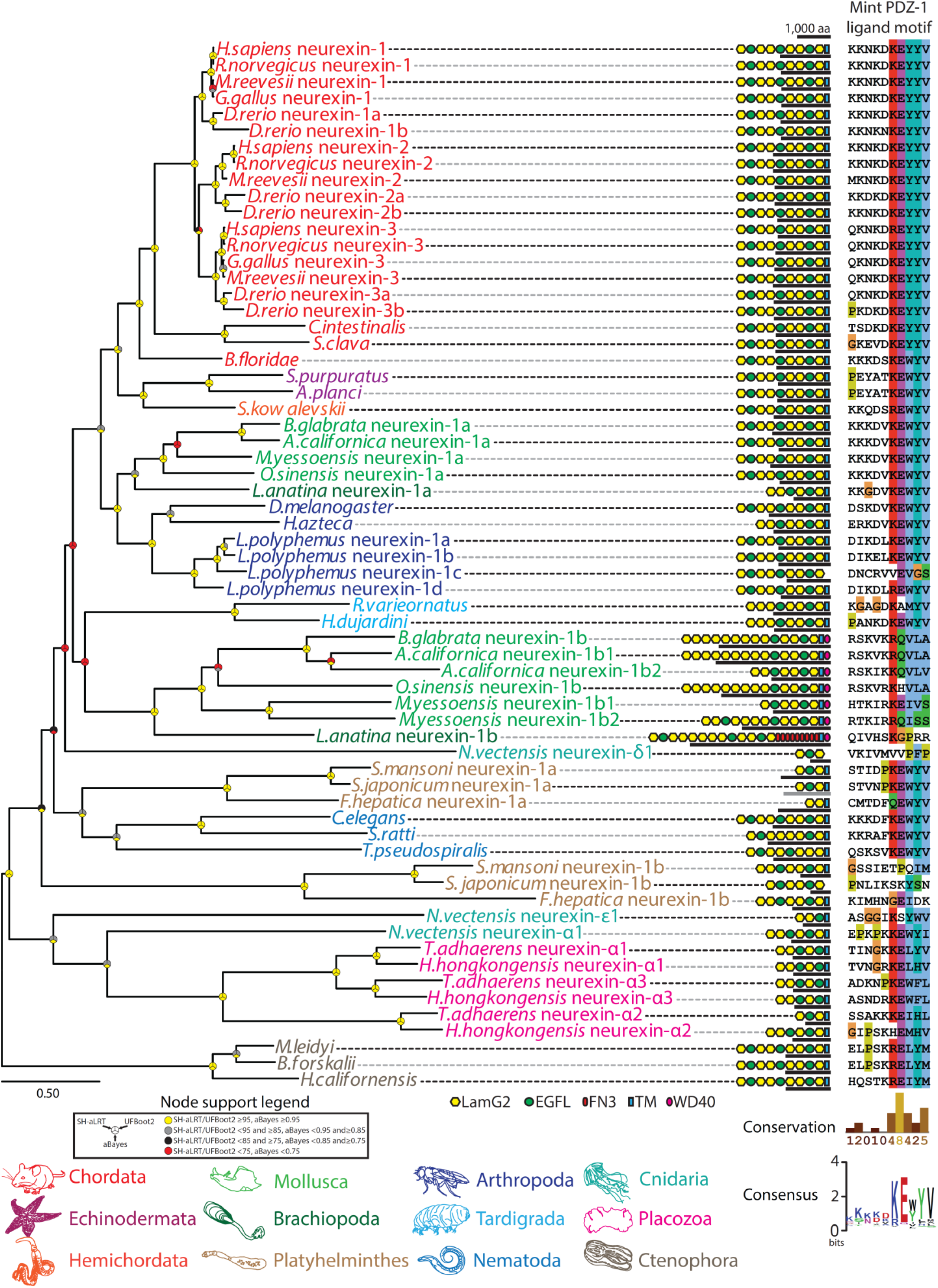
Maximum likelihood protein phylogeny of neurexin-1 homologues from various representative animals. Names of species and genes are colour-coded according to phylum. To the right of the phylogenetic tree is an alignment of the extreme C-terminal sequences of the proteins, illustrating deep conservation of the Mint PDZ-1 ligand motif. The lengths of the neurexin-1 homologues are indicated with black bars, and protein domains predicted with InterProScan are illustrated (LNS: laminin, neurexin, sex-hormone binding globulin domain; EGFL: epidermal growth factor-like domain; FN3: fibronectin type III domain; WD40: tryptophan-aspartic acid 40 repeat domain). Node support values generated with SH-aLRT, UFBoot2, and aBayes are indicated by symbols according to the provided legend.

Like PDZ-1, the Mint PDZ-2 domain mediates several distinct interactions from PDZ-1, including with the copper cofactor to the antioxidant superoxide dismutase-1 (CCS) [74] and the Ca^2+^-regulated transcription factor NK-κB [75] (Figure 1E). Lastly, we note unique tetra-trico-peptide repeat (TCP) protein-protein interaction domains upstream of the PTB domain in the two examined placozoan Mint orthologues, notable because placozoans exhibit an enrichment of this particular domain within their proteomes [76], and like PDZ domains, some TPR domains can recognize C-terminal ligands of target proteins [77].

### Stronger sequence divergence of Mint PDZ-1 vs. PDZ-2 suggests placozoans and ctenophores lack the capacity for a Mint-Ca_V_2 interaction

To better understand the conservation between orthologous Mint PDZ domains, we generated protein sequence alignments and predicted secondary structures of select PDZ-1 and PDZ-2 domains (Figure 4A and B). Both PDZ-1 and PDZ-2 bear a standard arrangement of beta strands (β1 to β6) and alpha helices (α1 and α2), with the exception of a short β5 strand, suggesting both adopt a mostly canonical PDZ domain tertiary structure [78]. For PDZ-1, we were able to map the positions of six amino acids whose sequence divergence correlates with evolutionary divergence in ligand specificity between different classes of PDZ domains [79]. These residues, which are clustered within and proximal to the β2 and β3 strands, are highly conserved within the PDZ-1 domains of bilaterian Mint orthologues, with some notable differences apparent for those from the non-bilaterian species *T. adhaerens* and *H. californiensis* (Figure 4A). More variable are five amino acids in the α2 helix that for the rat Mint-1 PDZ-1 domain were shown to exhibit strong spectral shifts in NMR spectrograms upon binding a Ca_V_2.2 channel DDWC ligand [48], with the exception of a cationic lysine/arginine residue in the tail end of α2 only absent in the *T. adhaerens* and *H. californiensis* orthologues.

**Figure 4.**
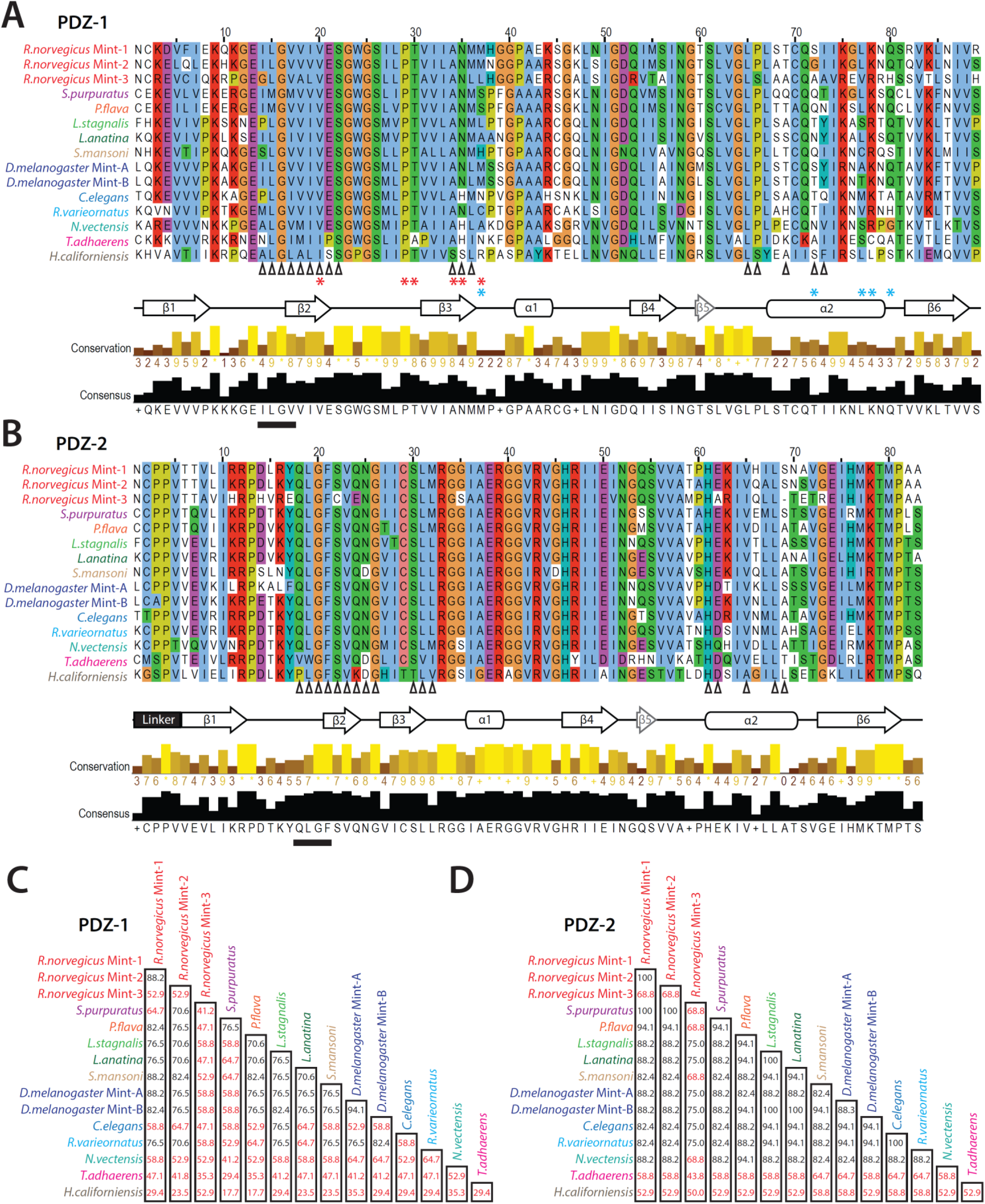
The Mint PDZ-1 domain is more divergent in protein sequence compared to PDZ-2. **(A)** Protein sequence alignment of select Mint PDZ-1 domains revealing deep sequence conservation throughout the secondary structure comprised of 6 β-strands and 3 α-helices. The black chevrons under the alignment indicate the position of the 17 amino acids correlated with ligand specificity [41], the red asterisks indicate positions that correlate with the evolution of distinct ligand preferences for PDZ domains in animals [79], and the blue asterisks denote amino acid positions that show major spectral shifts upon binding to the Ca_V_2 channel DDWC ligand in NMR studies [48]. **(B)** Protein sequence alignment of select Mint PDZ-2 domains. Like panel A, the black chevrons under the alignment indicate the position of the 17 amino acids correlated with ligand specificity [41]. For panels A and B, the black bars indicate the position of the ligand carboxylate accommodating loop within the PDZ domain ligand-binding groove, the latter comprised of the β2 strand and the α2 helix. The labelled arrows placed under the alignments denote expected locations of β strands, and the labelled bars the locations of alpha helices. **(C)** Sequence identity matrix (percent identity) of 17 aligned amino acids within the PDZ domain ligand-binding groove shown to correlate with ligand specificity [41]. Font colors for names of species/genes indicate of phylum according to the legend in Figure 1A. Percent identity values that fall below 70%, described as the threshold for correlated ligand specificity, are depicted in red font. **(D)** Similar sequence identity matrix as described in panel C, but for the Mint PDZ-2 domain.

Together, the noted sequences above overlap with 17 amino acid positions that correlate with distinct ligand specificities of PDZ domains determined via high-throughput peptide phage display screening [41]. Specifically, 17 amino acids clustered within the β2 and β3 strands and the α2 helix (Figure 4C), which mediate key contacts with C-terminal peptide ligands [80], were shown to strongly predict common ligand-binding specificity when conserved above 70% sequence identity [41]. A sequence identity matrix of these 17 aligned amino acids reveals stronger conservation in Mint PDZ-2 compared to PDZ-1 (Figure 4C and D). For PDZ-1, notable divergence is evident for rat Mint-3 and the Mint orthologues from *T. adhaerens* and *H. californiensis*, with all comparisons yielding alignment scores below 70%.

Sequence comparisons of PDZ-1 show an overall reduction in alignment scores compared to PDZ-2, particularly so for rat Mint-3 and orthologues from *Strongylocentrotus purpuratus*, *C. elegans*, *N. vectensis*, and *H. californiensis*. The non-bilaterian Mint PDZ-1 domains are highly divergent, both from each other and from bilaterian orthologues, producing alignment scores ranging between 64.7% and 17.7%. Notably, a correlation between global sequence identity of entire PDZ domains and ligand specificity was also reported, with a sequence identity cut-off of 50% [41]. Alignment matrices of entire Mint PDZ-1 and PDZ-2 domains reveal that most comparisons yield sequence identity scores above 50%, with only the PDZ-1 domains from the *T. adhaerens* and *H. californiensis* orthologues consistently falling below the 50% cut-off (Figure S1). Altogether, the examined sequence features suggest that Mint PDZ-1 ligand binding is more divergent compared to PDZ-2, particularly so for the Mint orthologues from placozoans and ctenophores which as noted above have Ca_V_2 channels lacking DDWC-like motifs (Figure 1A).

To date, the structural arrangement of the two tandem PDZ domains of Mint have not been experimentally determined. However, separate NMR structures of the human Mint-1 PDZ-1 and PDZ-2 domains are available (PDB numbers 1U37 and 1Y7N, respectively) [48, 81]. We therefore used AlphaFold2 [82] to predict structures of entire PDZ-1/PDZ-2/P1BM modules of Mint orthologues from *R. norvegicus* (Mint-1), *N. vectensis*, *T. adhaerens*, and *H. californiensis* (Figure 5A). All four structures were predicted with high to very high confidence, in particular the *H. californiensis* orthologue (Figure S2). Structural alignment between the solved Mint NMR structures and the predicted PDZ-1/PDZ-2/P1BM structure of *R. norvegicus* Mint-1 reveals root mean square deviation (r.m.s.d.) values of 0.87 and 0.67 angstroms for PDZ-1 and PDZ-2, respectively, indicating low deviation in spatial positioning of backbone alpha carbons between the solved and predicted structures. In all predicted structures, the extreme C-terminus of P1BM is found nestled within the ligand-binding groove of PDZ-1 comprised of the β2 strand and the α2 helix, consistent with NMR analyses indicating that the extreme C-terminus of P1BM inserts in a perpendicular orientation into this region of PDZ-1 leading to inhibition of ligand binding (*i.e.*, auto-inhibition) [48]. Interestingly, within all four structures the P1BM sequence bears a predicted alpha helix positioned between PDZ-1 and PDZ-2, corresponding to highly conserved residues including an arginine/lysine near the center of the alpha helix that is predicted to form a salt bridge with a conserved glutamate/aspartate residue 6 amino acids downstream (Figures 1D and 5B). The only orthologues lacking this cationic residue are from the ctenophore species *M. leidyi* and *H. californiensis*. Nonetheless, the *H. californica* Mint P1BM bears a polar asparagine residue at this position which is predicted to hydrogen bond with the conserved glutamate downstream. Clearly, future studies to determine whether these predicted structural elements serve to project and/or stabilize the extreme C-terminus of P1BM into the ligand-binding groove of PDZ-1 are warranted.

**Figure 5.**
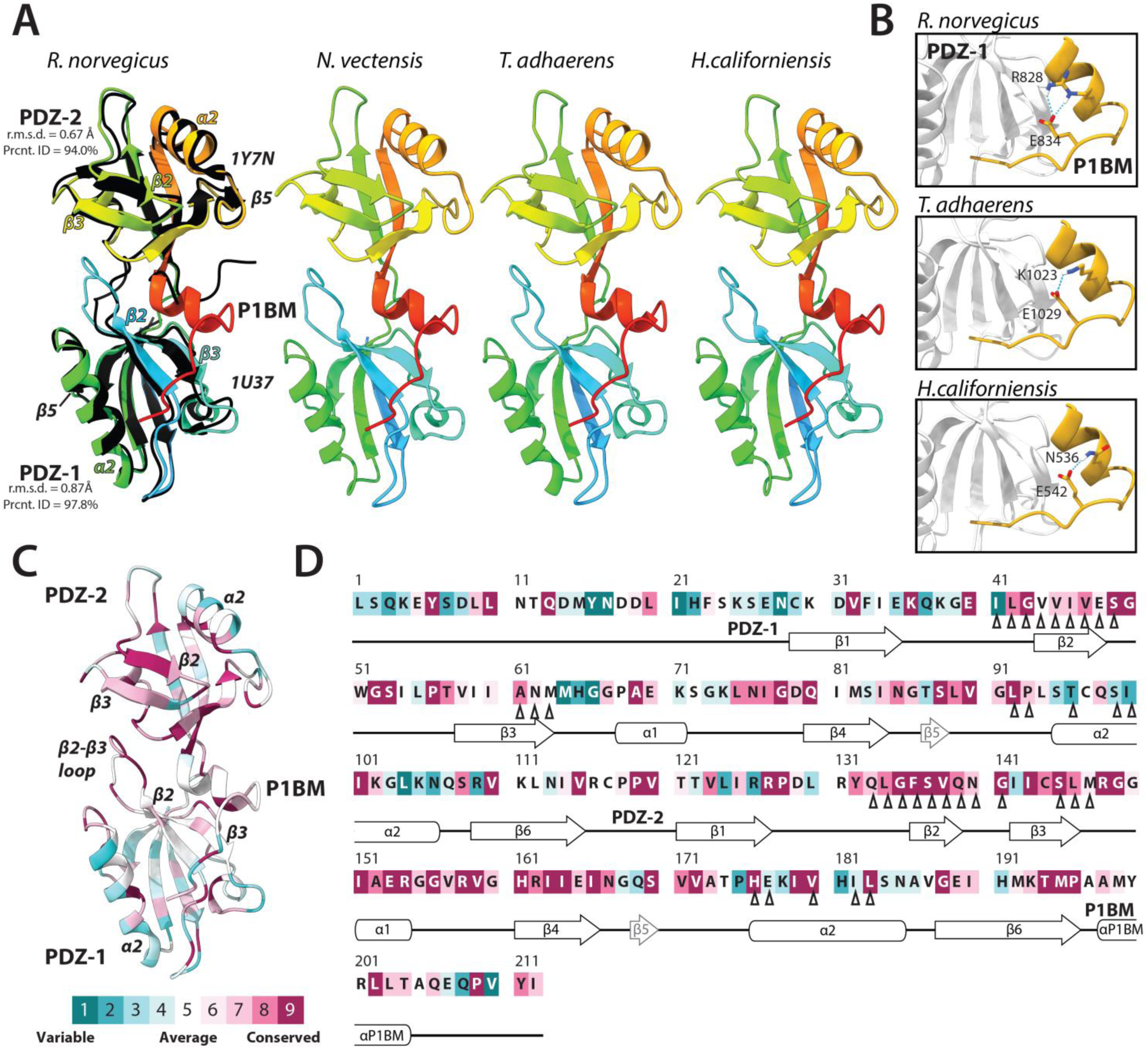
AlphaFold2-predicted structures of Mint PDZ-1, PDZ-2 and the auto-inhibitory P1BM element are consistent with the separately resolved structures of human Mint-1 PDZ-1 and PDZ-2. **(A)** Ribbon diagrams of AlphaFold2-predicted structures of the tandem PDZ domains of select Mint proteins, along with the C-terminal P1BM auto-inhibitory domain. Ribbon diagrams of the predicted structures are rainbow colored, with N-terminal elements in blue and the C-terminal in red. The black ribbon structures structurally aligned with the *Rattus norvegicus* AlphaFold2 structure are NMR structures of the human PDZ-1 and PDZ-2 domains, which were resolved as separate structures in two different studies (PDB accession numbers 1U37 and 1Y7N, respectively). Note that for all examined PDZ-1/PDZ-2/P1BM structures, AlphaFold2 predicts the existence of a novel alpha helix that helps orient the C-terminal auto-inhibitory peptide into the PDZ-1 ligand-binding groove. **(B)** Depiction of predicted intra-molecular hydrogen bonds between a conserved glutamate near the C-terminus of Mint and amino acids upstream within a novel predicted helix of the P1BM element. **(C)** Depiction of the *R. norvegicus* PDZ-1/PDZ-2/P1BM AlphaFold2 structure color-coded according to sequence conservation across phyla using ConSurf [83]. **(D)** Primary sequence of the *R. norvegicus* Mint-1 PDZ1-PDZ2-P1BM elements, color-coded according to the ConSurf sequence conservation analysis. The black chevrons under the alignment indicate the position of the 17 amino acids correlated with ligand specificity [41]. For panels B and C, purple denotes highly conserved sequences/structural regions, while blue denotes highly variable sequences/regions (legend at the bottom of panel E).

Next, we used the program ConSurf [83] to map sequence conservation of aligned PDZ domains (*i.e.*, the single representatives from each phylum in Figure 4A and B) onto the predicted PDZ-1/PDZ-2/P1BM structure of *R. norvegicus* Mint-1 (Figure 5C and D). As expected, PDZ-2 shows stronger conservation compared to PDZ-1 along the entire structure, particularly along the β2 strand and residues in the α2 helix that together comprise the ligand-binding groove. Also deeply conserved is a TMP amino acid motif at the C-terminal end of β6 which demarks the boundary between PDZ-2 and P1BM. In PDZ-1, strong conservation is apparent at the interface between the β1-β2 linker and the β2 strand, a region that harbours the carboxyl accommodating loop for bound C-terminal ligands. In addition, the linkers between β2-β3 and the α1-β4 are considerably conserved within PDZ-1, despite the former being predicted as disordered (Figure S2) [84].

### Mint and Ca_V_2c channels from the cnidarian sea anemone Nematostella vectensis interact in vitro and are co-expressed in dissociated neurons

We took advantage of the extensive whole animal transcriptome data available for *N. vectensis* in the NvERTx database [85] to explore the expression patterns of Mint, CASK, Veli, and its three Ca_V_2 channel isotypes through embryonic development, along with the transcription factors forkhead box protein L2 (FoxL2) and homeobox protein Otx1 which delineate two distinct populations of neurons in adult *N. vectensis* [86]. NvERTx also provides information on the developmental co-expression of genes, by partitioning genes with similar expression profiles into eight defined clusters [85]. During embryonic development, the Ca_V_2c channel, as well as CASK, OtxC, and FoxL2 exhibit general increases in mRNA expression, most markedly within the first 25 hours after fertilization (*i.e.*, the blastula stage; Figure 6A). This contrasts the mRNA expression of Ca_V_2a, Ca_V_2b, and Mint, which exhibit sharp decreases during the blastula stage, but become stabilized as embryos transition through the gastrula, planula, and juvenile stages. Notably, Ca_V_2a and Ca_V_2b are part of the same co-expression cluster (E-2), suggesting they are part of a common cellular gene regulatory network [85]. However, both cellular transcriptome analysis and mRNA *in situ* hybridization indicate these two genes are expressed in different cell types, with Ca_V_2a being restricted to cnidocytes (stinging cells), and Ca_V_2b being expressed more broadly in cells within the mouth and tentacles [2, 87]. Also notable is that Mint, CASK, and Veli, which as noted form a conserved tripartite complex in bilaterians, fall within different co-expression clusters, and that none of the Ca_V_2 channel genes share a cluster with Mint or the neuronal marker genes, but that Ca_V_2c and CASK do fall within a common cluster.

**Figure 6.**
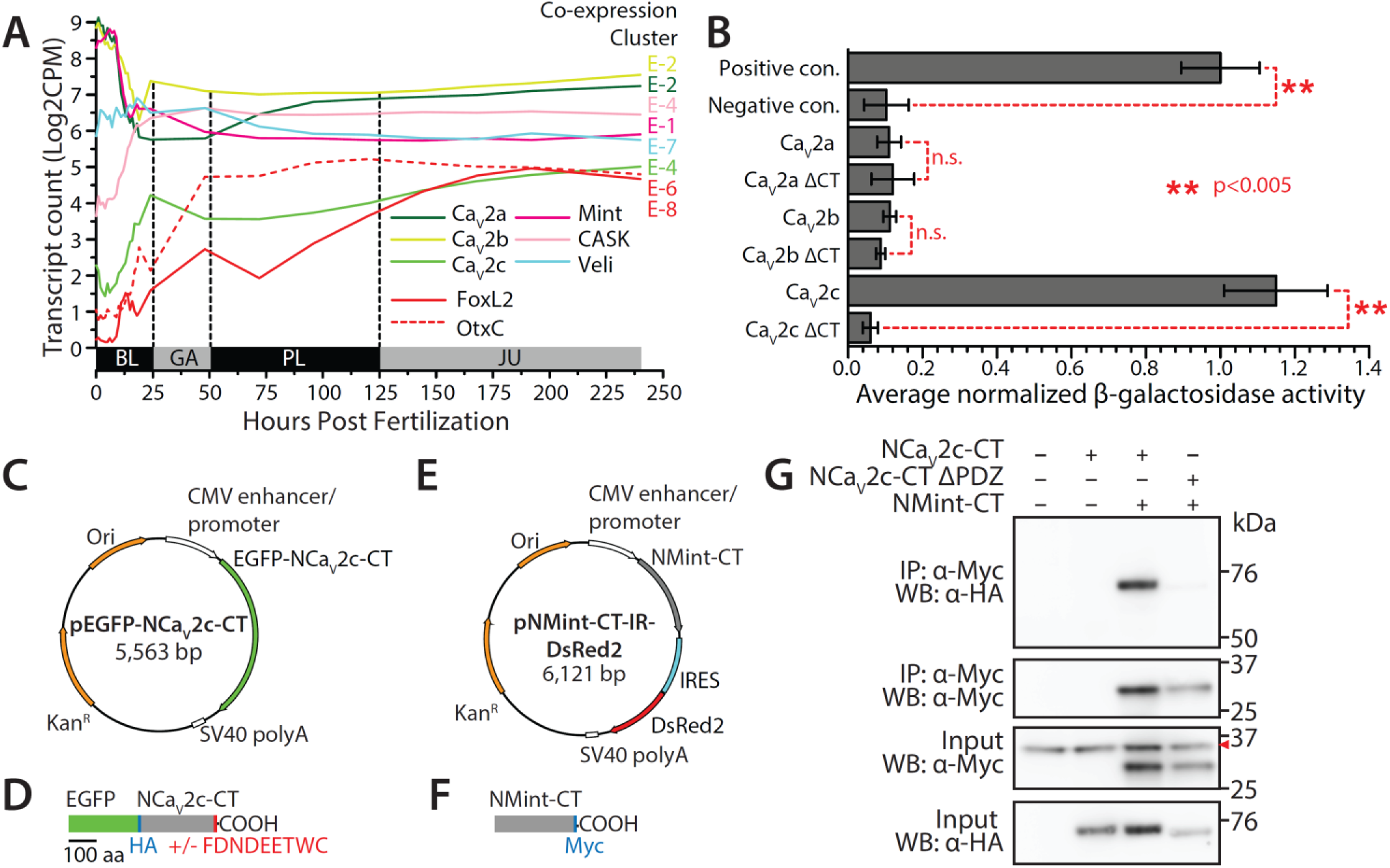
Mint and Ca_V_2c from the cnidarian *N. vectensis* interact *in vitro*. **(A)** Plot of mRNA transcript counts (Log2 of counts per million) of relevant gene transcripts in the developmental transcriptome *N. vectensis* obtained from the NvERTx database [85]. Included are the neuronal marker genes FoxL2 and OtxC which delineate two separate populations of neurons in the adult nervous system [86]. To the right of each line in the plot is an indication of the NvERTx co-expression cluster that each transcript is assigned to. BL = blastula, GA = gastrula, PL = planula, JU = juvenile. **(B)** Plot of average β-galactosidase activity (plus/minus standard deviation), normalized to a control positive interaction, measured for directed yeast 2-hybrid experiments between the *N. vectensis* Mint PDZ-1/PDZ-2/P1BM module and the Ca_V_2a to Ca_V_2c C-termini. The dashed red lines and asterisks denote statistical differences for two sample T-tests. **(C)** Map of the pEGFP-C1 plasmid construct use to express the *N. vectensis* Ca_V_2c channel C-terminus protein in HEK-293T cells as a HA-tagged GFP fusion protein. **(D)** Illustration of the *in vitro* expressed Ca_V_2c protein, depicting inclusion/exclusion of the 9 C-terminal amino acids bearing a putative Mint PDZ-1 domain ligand sequence. **(E)** Map of the pIRES2-DsRed2 plasmid construct use to express a portion of the *Nematostella* Mint PDZ1-PDZ2-P1BM protein with a Myc tag at the C-terminus. **(F)** Illustration of the *in vitro* expressed Mint protein. The lengths of the recombinant Ca_V_2c and Mint proteins are denoted by a shared scale bar of 100 amino acids. **(G)** Co-immunoprecipitation experiment showing a robust interaction between the *N. vectensis* Mint and the Ca_V_2c channel proteins *in vitro*. This interaction is completely disrupted upon deletion of the Ca_V_2c channel C-terminal FDNDEETWC_COOH_ sequence. The red chevron denotes non-specific labeling of immunoglobulins by the α-Myc antibodies.

Considering the observed sequence and predicted structural similarity between the *N. vectensis* Mint PDZ-1 domain with its bilaterian orthologues (Figures 4, S1, and 5), and the presence of conserved C-terminal DDWC-like motifs among all three cnidarian Ca_V_2c channel isotypes (Figure 1A), we reasoned that this protein-protein interaction might be conserved in cnidarians. To test this possibility, we conducted directed yeast 2-hybrid experiments to explore whether the C-terminal portion of *N. vectensis* Mint, containing the PDZ-1/PDZ-2/P1BM elements of the protein (*i.e.*, D1141 to I1350 of NCBI accession number XP_032220836.1; Supplementary File 1), could bind to the 70 most distal amino acids of the three corresponding *N. vectensis* Ca_V_2a, Ca_V_2b, and Ca_V_2c channel C-termini. Quantification of β-galactosidase activity, normalized to positive control values for known interacting proteins (murine p53 and the SV40 large T-antigen), revealed a strong interaction between the *N. vectensis* Mint PDZ-1/PDZ-2/P1BM protein and the 70 most distal amino acids of Ca_V_2c, but not Ca_V_2a or Ca_V_2b (Figure 6B). The interaction between Mint and Ca_V_2c was completely disrupted upon removal of the last four amino acids of the channel C-terminus (*i.e.*, ΔETWC_COOH_). We next sought to validate this finding by conducting co-immunoprecipitation experiments with recombinant, epitope-tagged proteins expressed in HEK-293T cells. Through gene synthesis, we cloned the entire C-terminus of the *N. vectensis* Ca_V_2c channel (*i.e.*, S1632 to C1896 of NCBI accession number XP_032220836.1; Supplementary File 1) into the mammalian expression vector pEGFP-C1, with and without the 9 distal C-terminal amino acids bearing the putative PDZ ligand sequence ETWC (Figure 6C). Expressed *in vitro*, this construct produces a chimeric protein with enhanced green fluorescent protein at its N-terminus (Figure 6D), which we found to enhance *in vitro* expression of the disordered channel C-terminus, plus a 9 amino acid haemagglutinin (HA) epitope tag used for the pulldown experiments. Similarly, we cloned the C-terminal portion of *N. vectensis* Mint, bearing the PDZ-1/PDZ-2/P1BM elements (E1076 to I1350 of NCBI accession number XP_032233969.1; Supplementary File 2), into the bicistronic expression vector pIRES2-DsRed2 (Figure 6E). This construct expresses red fluorescent protein separately from the recombinant Mint protein, the latter engineered to contain a 10 amino acid myc epitope tag at its C-terminus for the pulldown experiments (Figure 6F). Western blotting of the two recombinant Ca_V_2c channel proteins expressed in HEK-293T cells, using monoclonal anti-HA antibodies, produced bands of appropriate molecular weight for both the wildtype channel C-terminus (*i.e.*, 59.5 kDa), and the C-terminal deletion variant (*i.e.*, ΔPDZ-L; 58.3 kDa; Figure 6G). Similarly, Western blotting of the recombinant Mint protein with anti-myc antibodies revealed an appropriate band of ∼30.8 kDa. In three separate pull-down experiments of the Mint protein, we were able co-immunoprecipitate the wildtype Ca_V_2c channel C-terminus, but not the deletion variant lacking 9 C-terminal amino acids (Figure 6G). Thus, the PTB/PDZ-1/PDZ-2/P1BM module of the *N. vectensis* Mint can bind the Ca_V_2c channel C-terminus *in vitro*, but only in the presence of the extreme C-terminus bearing the ETWC motif. Altogether, these analyses indicate that the capacity for Mint to bind the Ca_V_2 channel C-terminus is conserved between cnidarians and bilaterians.

Next, we sought to determine whether the mRNAs of these two genes are co-expressed in dissociated neurons, along with the neuronal marker genes OtxC and FoxL2 [86]. Co-labelling of freshly dissociated cells from tentacle bud stage *N. vectensis* specimens with fluorescently labeled RNAscope probes against Ca_V_2c, FoxL2, and Mint (Figure 7A to C’) revealed that most cells that express Ca_V_2c also express Mint, and that roughly one fifth of labelled cells express Ca_V_2c, Mint, and FoxL2 together (18.1 ±1.6%) (Figure 7G). Separate experiments on cells co-labelled with probes against Ca_V_2c, OtxC, and Mint (Figure 7D to F’) revealed similar co-expression of Ca_V_2c and Mint (26.8 ±2.4% of 484 cells), and of Ca_V_2c, Mint, and OtxC together (12.9 ±1.4%) (Figure 7H). In both experiments co-expression of all three mRNA targets was observed in cells with morphology characteristic of unipolar neurons (Figure 7C, D, and E), bipolar neurons (Figure 7A and F), and tripolar neurons (Figure 7B), identified as distinct neuron subtypes in transgenic *N. vectensis* polyps [88]. Given that FoxL2 and OtxC are expected to label neurons, it appears as though the Ca_V_2c channel is expressed in neurons, together with Mint. Furthermore, Mint appears to have broader cellular expression than the other tested transcripts, since 23.9 ±4.3% and 26.9 ±1.9% of all cells in the two separate experiments were labeled only with the Mint probe (Figure 7G and H). Morphologically, some of Mint-positive cells resemble gland cells (*e.g.*, middle cell on Figure 7A or upper right cell on Figure 7E), shown to exhibit high diversity in *N. vectensis* polyps via single cell RNA-seq [86, 89].

**Figure 7.**
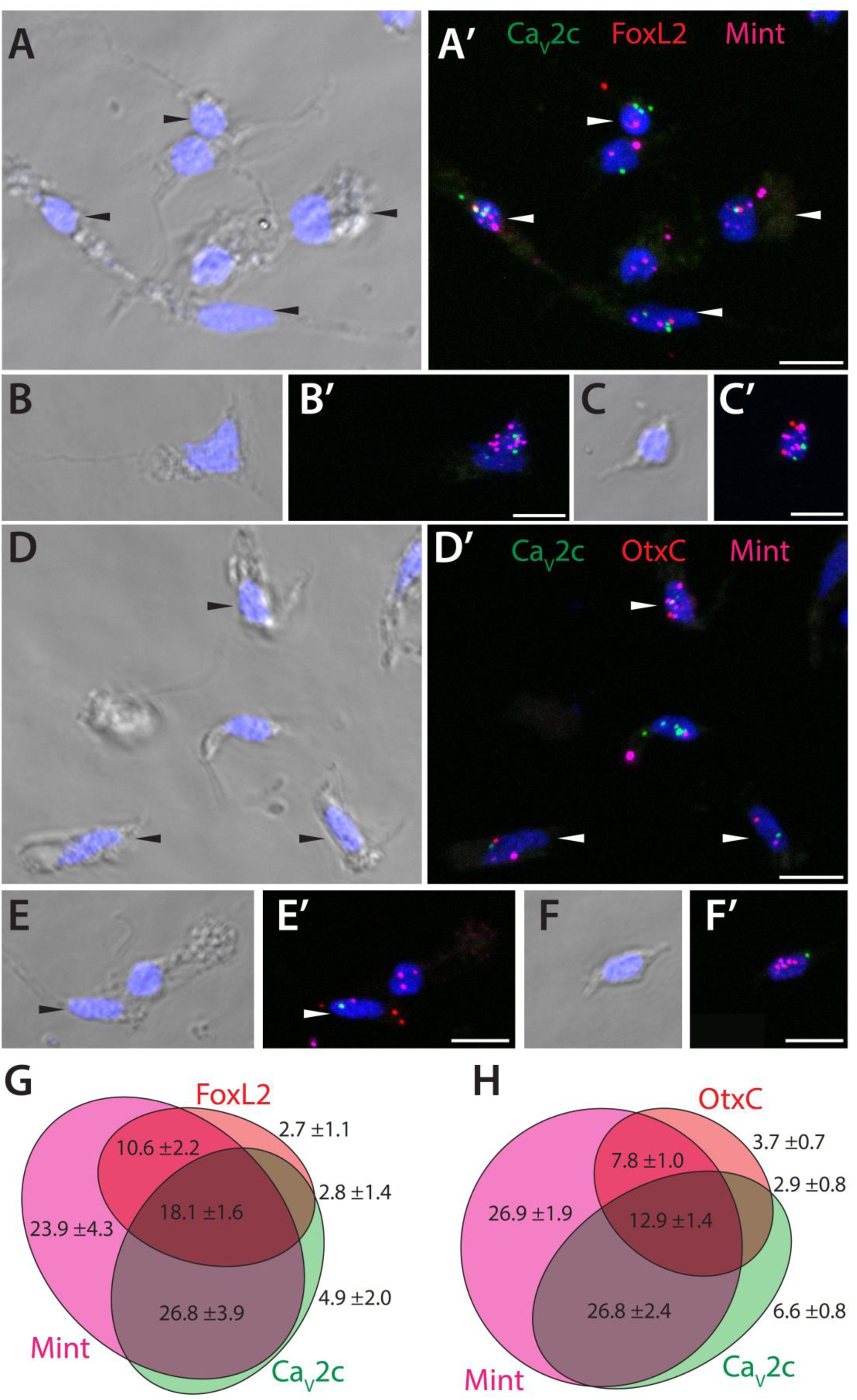
Mint and Ca_V_2c channels are co-expressed in dissociated *N. vectensis* neurons. **(A to C’)** Transmitted light (A to C) and fluorescent (A’ to C’) micrographs of dissociated *N. vectensis* cells subject to *in situ* hybridization with RNAscope probes for the Ca_V_2c channel (green), Mint (magenta), and FoxL2 (red). **(D to F’)** Transmitted light (D to F) and fluorescence (D’ to F’) micrographs of dissociated cells probed for the Ca_V_2c channel (green), Mint (magenta), and OtxC (red). White chevrons denote cells where all three mRNAs could be detected. White chevrons in panels B to G’ denote cells where all three mRNAs were detected as co-expressed. Scale bars represent 10 µm. **(G)** Venn diagram showing average single cell counts for co-expression of Mint, Ca_V_2c and FoxL2, showing that most cells expressing Ca_V_2c also express Mint, and that over 1/3 of cells expressing Ca_V_2c also express the neural marker FoxL2. **(H)** Venn diagram showing average single cell counts for co-expression of Mint, Ca_V_2c and OtxC, showing that most cells expressing Ca_V_2c also express Mint, and under 1/3 of cells expressing Ca_V_2c also express the neural marker OtxC.

To validate these observations, we mined the recently updated single cell transcriptome data available for *N. vectensis* [90]. Apparent is that Ca_V_2c mRNA expression is largely restricted to neurons and related secretory cells, as well as mucin-producing secretory gland cells (Figure 8A and B). Indeed, Ca_V_2c expression is much more prevalent in neuronal metacells comparted to Ca_V_2a and Ca_V_2b, present in 401 out of 3,709 neuronal metacells vs. only 57 for Ca_V_2a and 56 for Ca_V_2b. Mint also shows prominent expression in neurons (427 neuronal metacells) and secretory cells but is also expressed more broadly in other cell types including cnidocytes, gastrodermis, and ectoderm (Figure 8C). Consistent with their suggested classification as neuronal marker genes [86], FoxL2 and OtxC show enriched expression in neurons (459 and 295 metacells, respectively), but also moderate expression in a few other cell types (*e.g.*, FoxL2 in cnidocytes and retractor muscle, OtxC in gastrodermis and embryonic mesoderm; Figure 8D and E). Further analysis confirmed co-expression of Mint and Ca_V_2c in 72 metacells, and triple expression of Ca_V_2c, Mint, and either FoxL2 or OtxC in 13 metacells (Figure 8F and G). Thus, the single cell RNA-Seq data confirms neuronal co-expression of Mint and Ca_V_2c channels. Nonetheless, the low percentage of neuronal metacells co-expressing these two genes, which is considerably lower than what we observed in our *in situ* experiments (Figure 7), prompted us to explore whether the RNA-Seq data might underestimate real co-expression values. To do this, we explored the neuronal expression of the two *N. vectensis* Ca_V_β ancillary subunits, β1 and β2, which are critical components of Ca_V_1 and Ca_V_2 channel complexes required for proper protein folding, membrane expression, and function of the pore-forming Ca_V_ subunit [91]. Hence, there is an expectation that Ca_V_2c and either β1 or β2 should be highly co-expressed in neurons. Interestingly, 620 neuronal metacells express the β1 subunit (accession number NVE4481), whereas only 18 express β2 (NVE10585), indicating that the former is the predominant neural subunit. An examination of co-expression with Ca_V_2c identified only 76 metacells for β1, and 5 for β2. An examination of co-expression with Ca_V_2c identified only 76 metacells for β1, and 5 for β2. Hence, of the 401 neuronal metacells that express Ca_V_2c, 18.0% co-express Mint, 19.0% co-express β1, and only 1.2% co-express β2. Based on these numbers, it seems reasonable to infer that the co-expression value for Ca_V_2c and Mint extracted from the RNA-Seq data underestimates co-expression that occurs in neurons.

**Figure 8.**
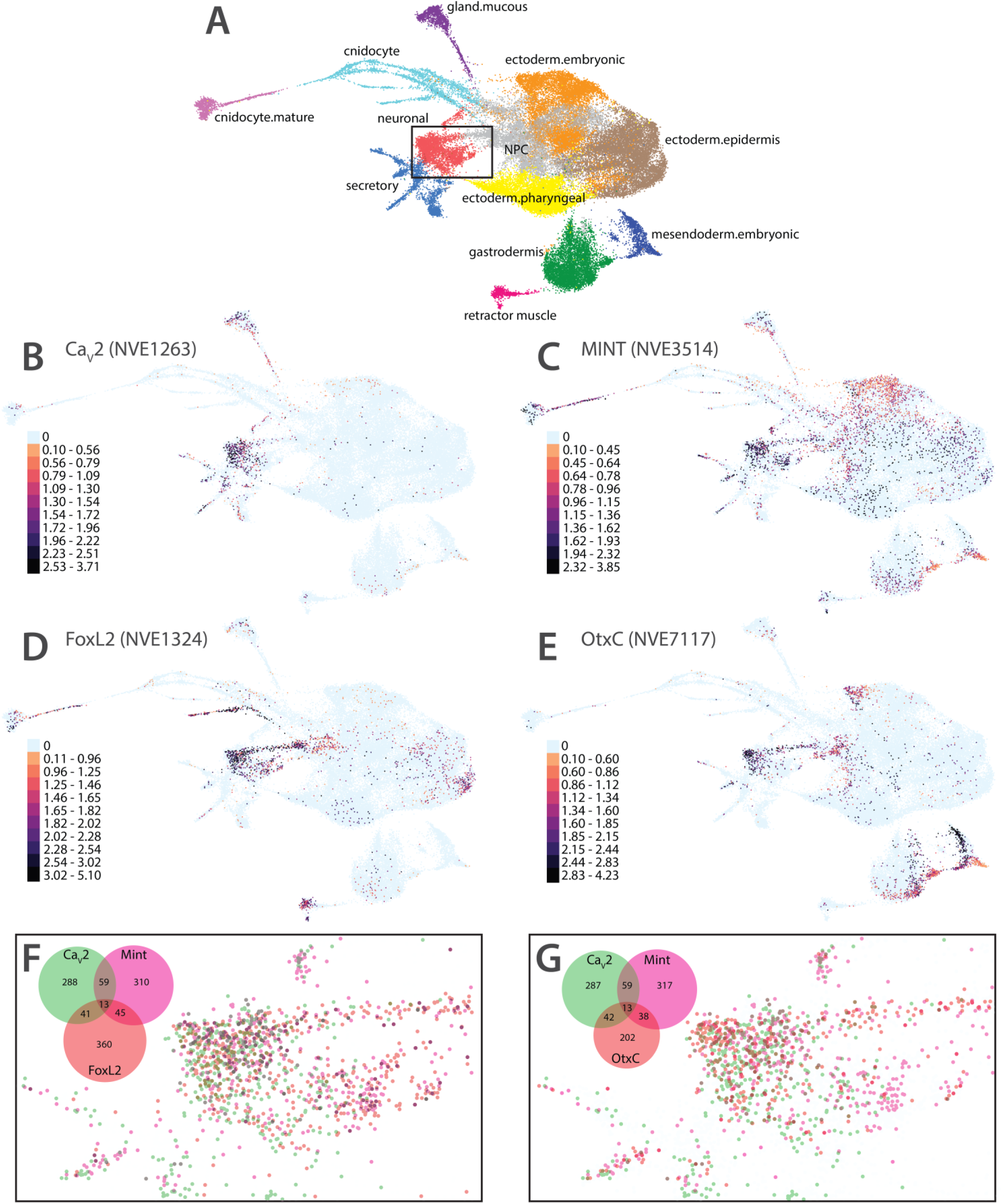
Mint and Ca_V_2c channels are co-expressed in *N. vectensis* neuronal metacells. **(A)** Complete cell state atlas of *N. vectensis* (from [90]). **(B)** Metacell expression of Ca_V_2c mRNA. **(C)** Metacell expression of Mint mRNA. **(D)** Metacell expression of FoxL2 mRNA. **(E)** Metacell expression of OtxC mRNA. **(F)** Overlay of metacells expressing Ca_V_2c, Mint, and FoxL2 within the neuronal cluster corresponding to the box in panel A. **(G)** Overlay of metacells expressing Ca_V_2c, Mint, and OtxC within the neuronal cluster corresponding to the box in panel A. For panel A, different colors are used to indicate different metacell clusters. For panels B to E, accession numbers for each gene are listed in brackets on the right of each gene name, and the legends within each panel are used to indicate number of cells within a given metacell. The Venn diagrams in the lower right of panels F and G illustrate the color scheme for co-expression within the different metacells.

### Mint PDZ domains from placozoans and ctenophores interact with Ca_V_2 channel C-termini in vitro

Placozoans like *T. adhaerens* are the most early diverging animals that possess all three types of metazoan Ca_V_ channel types, Ca_V_1 to Ca_V_3 [2, 4, 92]. Previously, the three *T. adhaerens* Ca_V_ channels, TCa_V_1 to TCa_V_3, were shown to conduct voltage-gated calcium currents *in vitro* with properties like those of their bilaterian orthologues [20, 30, 93].

Nonetheless, the absence of a DDWC-like C-terminus in placozoan Ca_V_2 channels (Figure 1A) suggests that the Ca_V_2-Mint interaction is not conserved in these animals. Alternatively, given the strong sequence divergence of the *T. adhaerens* Mint PDZ domains (Figures 2C and S1), the interaction may occur but through unique ligand binding properties of PDZ-1. To explore these possibilities, we sought to emulate the *in vitro* co-immunoprecipitation experiments done for *N. vectensis* using the corresponding *T. adhaerens* proteins. Thus, we similarly cloned the C-terminus of TCa_V_2 *(i.e.*, R1646 to V2093 of NCBI accession number QOQ37977.1; Supplementary File 1) into the pEGFP-C1 plasmid and tagged it with EGFP and an HA epitope at its N-terminus, and the PTB/PDZ-1/PDZ-2/P1BM region of *T. adhaerens* Mint (K760 to I1034 of NCBI accession number GHJI01000994.1; Supplementary File 2) into pIRES2-DsRed2 tagged with a C-terminal myc tag (Figure S3A and B). Unfortunately, the recombinant TCa_V_2 protein proved difficult to express in HEK-293T cells (Figure S3C), perhaps due to its extended length of 448 amino acids compared to only 265 for *N. vectensis* Ca_V_2c. As a result, an interaction between Mint and TCa_V_2 could neither be observed nor ruled out via co-immunoprecipitation.

As an alternative approach, we sought to identify putative binding partners of the *T. adhaerens* Mint PDZ-1/PDZ-2/P1BM protein, as well as PDZ-1 by itself, by generating and screening a custom yeast 2-hybrid library derived from *T. adhaerens* whole animal mRNA. Searching this library with PDZ-1/PDZ-2/P1BM as bait (*i.e.*, K825 to I1034) identified 12 putative binding partners, all but three bearing a valine residue at the extreme C-terminus (Table 1A). Two of the identified prey proteins, homologues of cathepsin D and a G protein γ subunit, produced 4 and 2 hits respectively, the latter notable in bearing a C-terminal sequence strikingly similar to that of TCa_V_2 (*i.e.*, SRCTLV_COOH_ vs. SKCTAV_COOH_). Using just Mint PDZ-1 as bait identified a total of 32 putative interactions (Table 1B), 17 of which corresponded to an artificial peptide encoded within the 3’ UTR of a betaine-homocysteine S-methyltransferase 1 homologue. Interestingly, this non-biological peptide has a C-terminal sequence ending with tyrosine and isoleucine (*i.e.*, YI_COOH_), resembling the Mint P1BM C-terminus of ETPEYI_COOH_ which based on homology and structural prediction is expected to insert into the PDZ-1 ligand-binding groove (Figures 1D, 5, and S2). Two additional prey sequences pulled from the cDNA library also had C-termini with terminal isoleucine residues, altogether suggesting that the PDZ-1 domain of *T. adhaerens* Mint has a high affinity for peptides bearing isoleucine in the last position. Although we did not directly test the interaction between the *T. adhaerens* Mint PDZ-1 domain to the downstream P1BM element, the strong binding of this PDZ-1 to P1BM-like C-terminal peptides suggests that auto-inhibition is an ancient and deeply conserved trait, also suggested by the deep sequence conservation of the P1BM element (Figure 1D). The second-most prominent identified prey sequences had C-termini ending with WV_COOH_ (6 hits), suggesting that like the PDZ-1 domain of mammalian Mint-1, the *T. adhaerens* orthologue has the capacity to bind C-terminal ligands with a tryptophan at position -1, coupled to a valine at position 0.

**Table 1.**
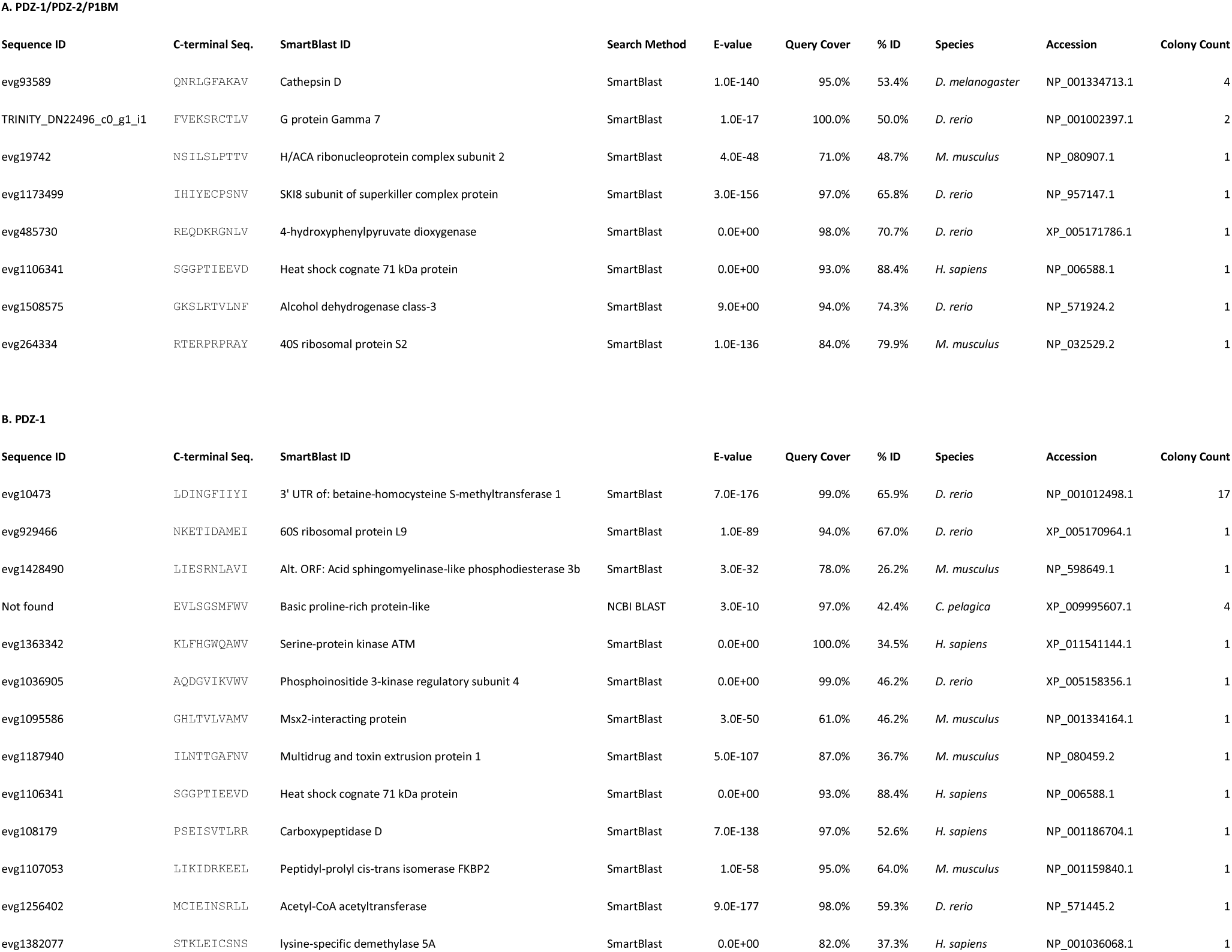
Annotation of *T. adhaerens* proteins found to interact with (A) the Mint PDZ-1/PDZ-2/P1BM and (B) the Mint PDZ-1 proteins in yeast 2-hybrid screens.

Intrigued by the similarity between the putative PDZ ligand sequence of the identified Gγ protein (SRCTLV_COOH_) and the TCa_V_2 channel (SKCTAV_COOH_), we next explored whether *T. adhaerens* Mint can bind the 70 most distal amino acids of the channel in directed yeast 2-hybrid experiments as we did previously for *N. vectensis* (Figure 6B). These experiments revealed a moderate interaction between the *T. adhaerens* Mint PDZ-1/PDZ-2/P1BM protein and the 70 most distal amino acids of TCa_V_2 (Figure 9), which was completely disrupted upon removal of the last four amino acids of the channel C-terminus (*i.e.*, ΔCTAV_COOH_). Instead, experiments using just PDZ-1 domain of *T. adhaerens* Mint failed to show an interaction with the TCa_V_2 C-terminus. Since our yeast 2-hybrid screen with PDZ-1 on its own identified putative ligands with a tryptophan residue in the -1 position (Table 1), we reasoned that this PDZ domain might share conserved binding properties with mammalian Mint PDZ-1 in this regard. Thus, we expanded these experiments to test *in vitro* interactions between *T. adhaerens* Mint and the Ca_V_2c channel from *N. vectensis* bearing the C-terminal sequence ETWC_COOH_. This revealed a moderate but statistically significant interaction (Figure 9). Interestingly, we did not detect an interaction between the complete *T. adhaerens* Mint PDZ-1/PDZ-2/P1BM module at the ETWC_COOH_ ligand, perhaps because the auto-inhibitory sequence of P1BM binds PDZ-1 to restrict its ability to bind the *N. vectensis* C-terminus (Figure 9).

**Figure 9.**
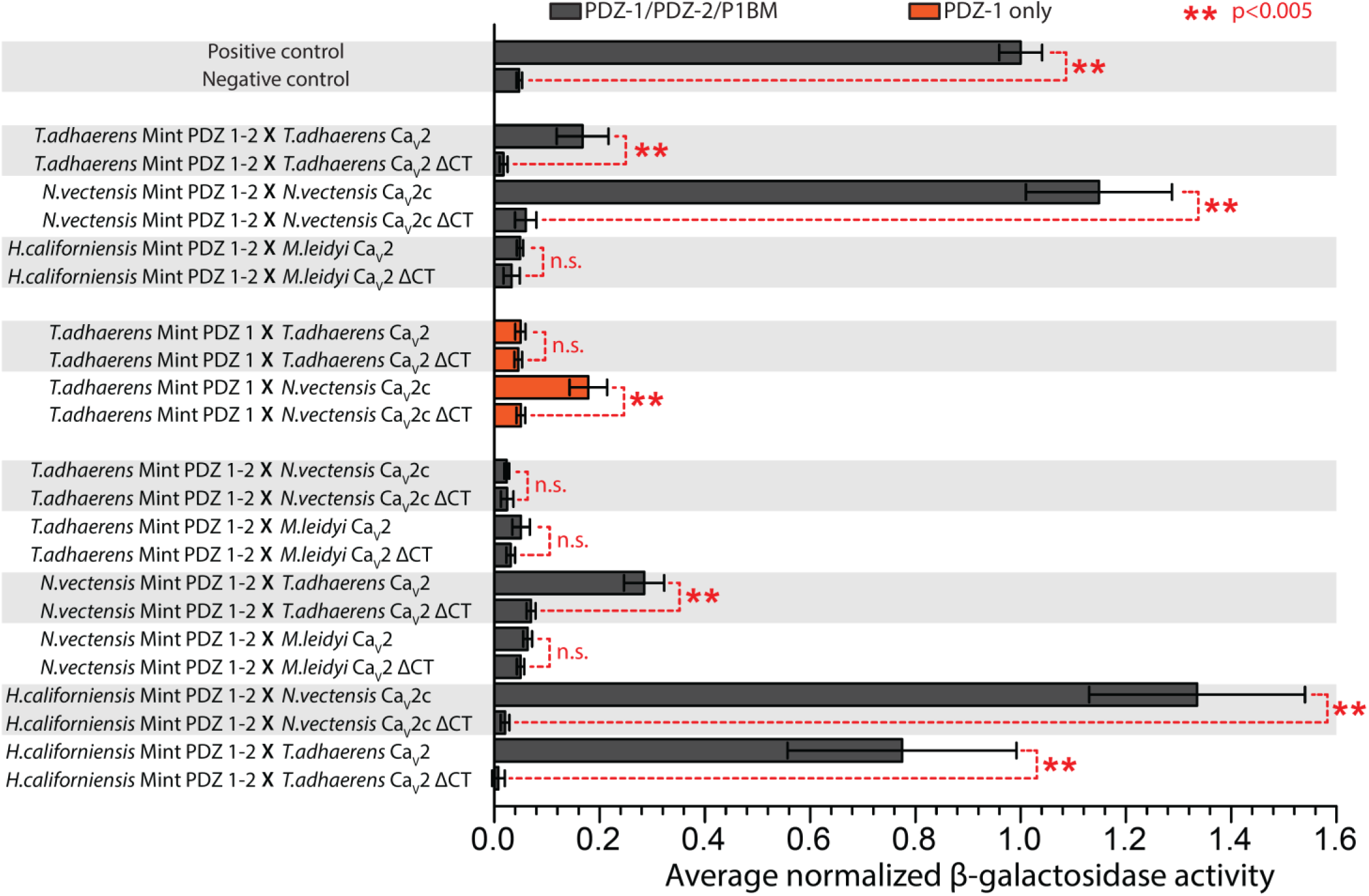
Yeast 2-hybrid analyses reveal deep conservation of the Mint-Ca_V_2 interaction. Plot of average β-galactosidase activity (plus/minus standard deviation), normalized to a control positive interaction, measured for various directed yeast 2-hybrid experiments. The dashed red lines and asterisks denote statistical differences for two sample T-tests.

We also tested *in vitro* interactions between ctenophore Mint and Ca_V_2 from the species *H. californiensis* (Mint) and *M. leidyi* (Ca_V_2) which have highly divergent Mint PDZ domains (Figures 4 and S1) and Ca_V_2 channel C-terminal sequences that are distinct from those of placozoans and cnidarians (Figure 1A). These experiments failed to detect an interaction between *H. californiensis* Mint (V344 to L547 of evg127856; Supplementary File 2) and the *M. leidyi* Ca_V_2 C-terminus (QOY24596.1; Supplementary File 1). Furthermore, neither the *T. adhaerens* nor the *N. vectensis* PDZ-1/PDZ-2/P1BM modules bound the *M. leidyi* Ca_V_2 channel C-terminal motif of SGDI_COOH_. To gain some insights into the binding properties of ctenophore Mint, we conducted cross-species experiments, finding strong interactions between the *H. californiensis* Mint PDZ-1/PDZ-2/P1BM and both the ETWC_COOH_ ligand of *N. vectensis* Ca_V_2c, and the CTAV_COOH_ ligand of *T. adhaerens* Ca_V_2 (Figure 9). The ability of ctenophore Mint to bind the distinct C-terminal ligands of placozoan and cnidarian Ca_V_2 channels prompted us to test whether *N. vectensis* Mint, like placozoan and ctenophore Mint, has dual binding capabilities in being able to bind the placozoan CTAV_COOH_ ligand in addition to its putative cognate ligand of ETWC_COOH_. These experiments revealed a moderate interaction between *N. vectensis* PDZ-1/PDZ-2/P1BM and the TCa_V_2 channel C-terminus (Figure 9).

Together, these various experiments reveal that *T. adhaerens* Mint can bind the TCa_V_2 channel C-terminus *in vitro*, albeit in a non-canonical manner as the binding is not mediated by PDZ-1 on its own. Like PDZ-1 in mammals, *T. adhaerens* Mint PDZ-1 can bind peptides with a tryptophan in the -1 position. Furthermore, the experiments demonstrate that cnidarian, placozoan, and ctenophore Mint PDZ domains can all recognize the placozoan Ca_V_2 ligand of CTAV_COOH_, but that none of the Mint PDZ-1/PDZ-2/P1BM modules from these animals bind the ctenophore Ca_V_2 channel C-terminus *in vitro*.

### Bacterial 2-hybrid screening uncovers unique binding properties of the T. adhaerens Mint tandem PDZ domains

Next, we sought to characterize the preferred ligand specificities of Mint PDZ domains from *T .adhaerens*, *N. vectensis*, and *H. californiensis*, as well as the brown rat *R. norvegicus*, by conducting bacterial 2-hybrid assays [94]. For each species, the protein-coding sequences of the PDZ-1/PDZ-2/P1BM module or PDZ-1 alone were cloned into the bacterial 2-hybrid vector pGHUC-ω-Erbin. These constructs were then screened for interactions with a library of random hexameric C-terminal peptides fused to a DNA-binding element in vector pB1H2-UV5-Zif268, permitting identification of ligand peptides *en masse* via Illumina sequencing [94]. Consistent with the ability of the rat Mint-1 PDZ-1 domain to bind the DDWC and DHWC sequences of Ca_V_2.1 and Ca_V_2.2 channels [46, 47], bacterial 2-hybrid screening with this domain identified a preferred ligand sequence of E/DWL_COOH_, resembling that of Ca_V_2 channels with an invariant tryptophan residue at position -1 preceded by a negatively-charged glutamate (E) or aspartate (D) at position -2 (Figure 10A).

**Figure 10.**
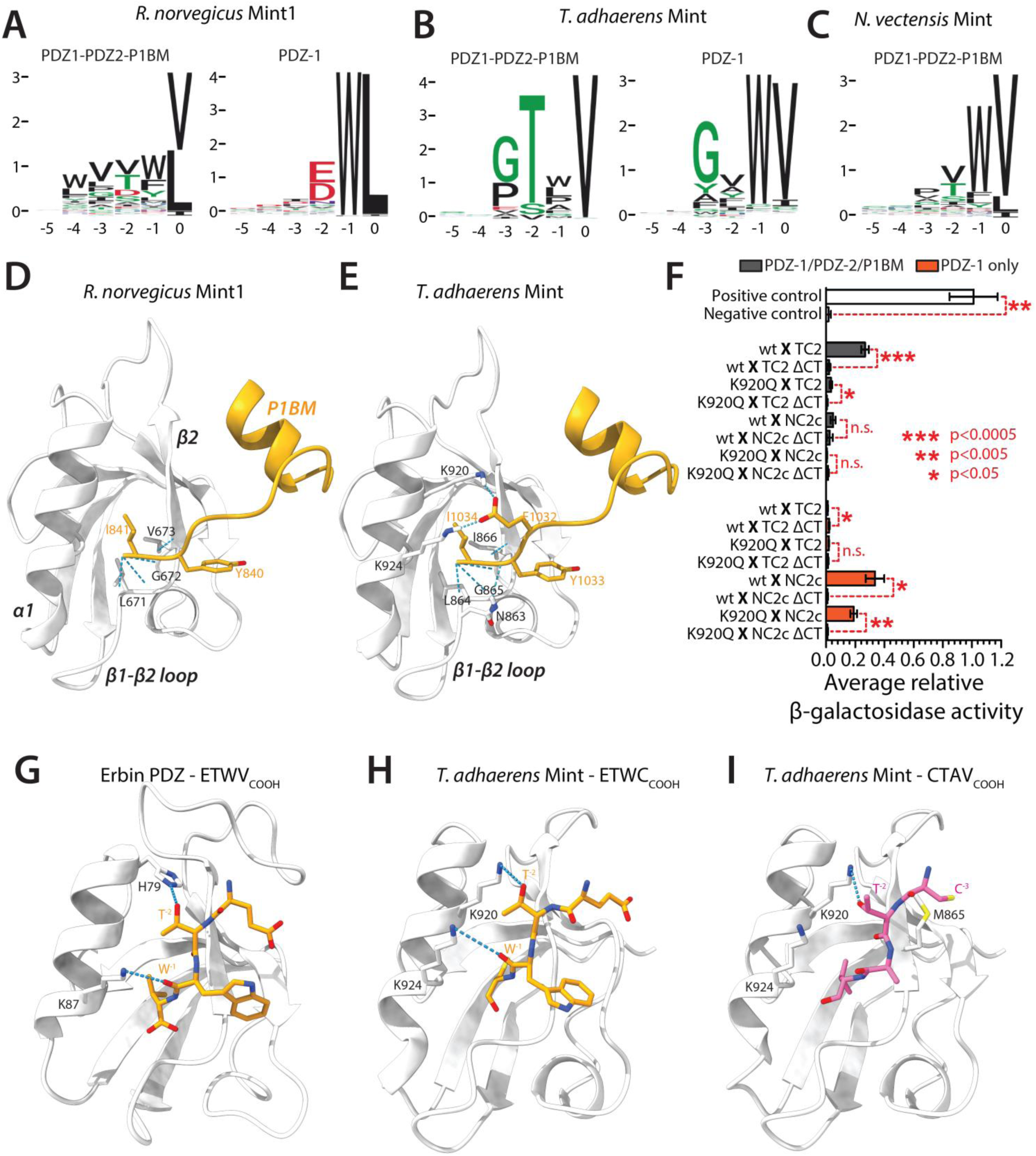
Bacterial 2-hybrid experiments, structural modelling, and mutagenesis shed light the unique ligand-binding properties of *T. adhaerens* Mint. **(A)** Sequence logos for the preferred ligand signatures of the *R. norvegicus* Mint PDZ-1/PDZ-2/P1BM protein (left), and the PDZ-1 domain by itself (right), determined via bacterial 2-hybrid screening. **(B)** Sequence logos for the preferred binding signatures of the *T. adhaerens* Mint PDZ-1/PDZ-2/P1BM protein (left), and the PDZ-1 domain by itself (right), determined via bacterial 2-hybrid screening. **(C)** Sequence logo for the preferred ligand signature of the *N. vectensis* Mint PDZ-1/PDZ-2/P1BM protein determined via bacterial 2-hybrid screening. **(D)** AlphaFold2 model of the *R. norvegicus* Mint-1 auto-inhibitory peptide of the P1BM element embedded within the PDZ-1 ligand-binding groove predicting. Weak Van der Waals interactions between the C-terminal isoleucine of P1BM and several amino acids within the PDZ-1 β1-β2 loop and β2 stand that would serve to stabilize the interaction are shown. **(E)** AlphaFold2 model of the *T. adhaerens* Mint auto-inhibitory peptide-PDZ-1 interaction, revealing unique hydrogen bonds between a glutamate residue two amino acids upstream of the C-terminal isoleucine (E_1034_) and two lysine residues in the α2 helix pf PDZ-1 (K_920_ and K_904_). **(F)** Plot of average β-galactosidase activity (plus/minus standard deviation), normalized to a control positive interaction measured for various directed yeast 2-hybrid experiments. The dashed red lines and asterisks denote statistical differences for paired T-tests (Legend: wt = wildtype *T. adhaerens* PDZ-1/PDZ-2/P1BM or PDZ-1; TC2 = TCa_V_2; NC2c = *N. vectensis* Ca_V_2c; ΔCT = last four amino acids removed). **(G)** Solved x-ray crystal structure of the human erbin PDZ domain bound to its preferred peptide ligand ETWV_COOH_ (PDB number 6Q0N). **(H)** Modelled structure of the *T. adhaerens* Mint PDZ-1 domain bound to its recognized ligand ETWC_COOH_ from the C-terminus of the *N. vectensis* Ca_V_2c channel. **(I)** Modelled structure of the *T. adhaerens* Mint PDZ-1 domain bound to the non-recognized ligand CTAV_COOH_ from the C-terminus of the *T. adhaerens* Ca_V_2 channel.

Notable however is the presence of a leucine at position 0 of the consensus ligand sequence, contrasting the cysteine residue conserved in Ca_V_2 channels (Figure 1A). Screening the entire PDZ-1/PDZ-2/P1BM module of rat Mint-1 revealed a markedly different ligand specificity, with a reduced selection for tryptophan at position -1, consistent with auto-inhibition of PDZ-1 by P1BM, as well as a slight enrichment for hydrophobic residues at positions -2 to -4, and a reduced preference for leucine at position 0 in lieu of the also hydrophobic residue valine.

For the *T. adhaerens* Mint PDZ-1 domain, the screening identified a preferred ligand sequence of GxWV_COOH_ (Figure 10B), which is consistent with the observed interaction of this protein with the *N. vectensis* Ca_V_2c channel C-terminus in our directed yeast 2-hybrid experiments (Figure 9), and C-terminal sequences of several interacting proteins identified in the yeast 2-hybrid screen (Table 1). Also consistent, the *T. adhaerens* PDZ-1/PDZ-2/P1BM module produced a ligand specificity of GTxV_COOH_, corroborating its ability to bind the TCa_V_2 channel C-terminus in directed yeast 2-hybrid experiments (Figure 9), as well as other proteins bearing TxV_COOH_ or V_COOH_ C-termini in the yeast 2-hybrid screening (Table 1). Notably, like rat Mint-1, the *T. adhaerens* Mint PDZ-1/PDZ-2/P1BM protein exhibits a diminished preference for tryptophan in the second last position, consistent with our inability to detect an interaction between this protein and the *N. vectensis* Ca_V_2c channel C-terminus in the yeast 2-hybrid assays (Figure 9), and with the occurrence of auto-inhibition of PDZ-1 by P1BM. A notable and unique feature of the *T. adhaerens* Mint PDZ-1/PDZ-2/P1BM compared to the rat and *N. vectensis* orthologues is a strong preference for threonine at position -2, and a common preference of PDZ-1/PDZ-2/P1BM and PDZ-1 alone to bind glycine and valine at positions -3 and 0, respectively (Figure 10A). As such, the most notable difference in ligand selectivity between the two *T. adhaerens* proteins are a tryptophan at position -1 for PDZ-1, instead of a threonine at position -2 for PDZ-1/PDZ-2/P1BM.

Unfortunately, neither of the two *H. californinesis* PDZ domain constructs produced viable ligand specificity signatures, nor did the *N. vectensis* PDZ-1 domain. However, we were able to generate a selectivity signature of WV_COOH_ for the *N. vectensis* Mint PDZ-1/PDZ-2/P1BM protein, which contrasts the signatures for rat and *T. adhaerens* orthologues by bearing a moderate preference for tryptophan at position -1 and corroborates the co-immunoprecipitation and yeast 2-hybrid experiments where we detected robust *in vitro* interactions of this protein with the ETWC ligand sequence of Ca_V_2c.

Indirectly, our analyses suggest that auto-inhibition of PDZ-1 by the P1BM module is conserved in the *T. adhaerens* Mint orthologue. To gain more insights into this possibility, we examined the AlphaFold2 structures of the rat and *T. adhaerens* PDZ-1/PDZ-2/P1BM modules, focusing on the predicted interaction between PDZ-1 and P1BM (Figure 10D and E). For both proteins, the P1BM element was predicted to insert the C-terminal YI_COOH_ motif into the PDZ-1 ligand-binding groove in a perpendicular orientation compared to the canonical orientation of PDZ domain C-terminal ligands (Figure 10D and E). For the rat orthologue, the C-terminal isoleucine of P1BM (I_841_) forms predicted hydrogen bonds with the residues L_671_, G_672_, and V_673_ comprising the highly conserved ligand carboxylate-binding loop in PDZ-1 (Figure 10D). This predicted arrangement is highly consistent with previous NMR studies which revealed a similar orientation of the C-terminal element of P1BM within the PDZ-1 ligand-binding groove, with hydrogen bonding between I_841_ and the residues L_671_ and G_672_ [48].

For the *T. adhaerens* protein, homologous interactions are predicted for the C-terminal residue I_1034_ of P1BM, which in the model forms hydrogen bonds with the carboxylate-binding loop residues L_864_, G_865_, and I_866_ of PDZ-1 (Figure 10E). However, there are also some additional hydrogen bonds predicted between Y_1033_ of P1BM with N_863_, as well as stronger salt bridges between an upstream glutamate in P1BM (E_1032_) and a set of lysine residues in the PDZ-1 domain α1 helix (K_920_ and K_924_). Interestingly, whereas the basic K_924_ residue is deeply conserved in the PDZ-1 domains of Mint, the upstream residue K_920_ appears to be a unique feature of *T. adhaerens* Mint, with most other orthologues possessing a glutamine in this position (Figure 4A). We therefore decided to explore whether this unique residue contributes to the unique ligand-binding properties of *T. adhaerens* Mint *in vitro*. Specifically, we repeated the directed yeast 2-hybrid experiments to test interactions between the *T. adhaerens* Mint PDZ-1/PDZ-2/P1BM and PDZ-1 proteins, and the TCa_V_2 and *N. vectensis* Ca_V_2c channel C-termini, comparing wildtype Mint variants to ones bearing a K_920_Q mutation within PDZ-1. Interestingly, this amino acid substitution completely disrupted the interaction between the *T. adhaerens* Mint PDZ-1/PDZ-2/P1BM protein and the TCa_V_2 channel C-terminal motif of CTAV_COOH_, while only reducing the interaction between PDZ-1 alone and the ETWC_COOH_ motif of *N. vectensis* Ca_V_2c (Figure 10F). Thus, while the unique K_920_ residue in PDZ-1 of *T. adhaerens* Mint does not seem critical for PDZ-1 binding to DDWC-like ligands, it plays a major role in the ability of PDZ-1/PDZ-2/P1BM to recognize and bind the TCa_V_2 CTAV_COOH_ ligand.

Lastly, to further explore why the *T. adhaerens* Mint PDZ-1 domain is able bind the *N .vectensis* Ca_V_2c peptide of ETWC_COOH_, but not its cognate Ca_V_2 peptide of CTAV_COOH_, we modelled these interactions for comparison with the solved X-ray crystal structure of the erbin PDZ domain bound to its preferred (and similar) ligand peptide ETWV_COOH_ [95] (PDB number 6Q0N). Interestingly, in both the resolved structure of erbin-ETWV_COOH_ and the modelled structure of Mint PDZ-1-ETWC_COOH_ (Figure 10G to I), a lysine residue in the distal end of the PDZ α1 helix (*i.e.*, K_87_ in erbin and the conserved K_924_ residue in Mint) hydrogen bonds with the backbone carbonyl oxygen of the peptide tryptophan, W^-1^, orienting its aromatic side chain over the proximal end of the β2 strand (Figure 10G and H). Given the deep conservation of an acidic residue in this position of Mint PDZ-1 (Figure 4A), including in all orthologues from species where an interaction with Ca_V_2 channel C-termini bearing W^-1^ residues has been confirmed *in vitro* (Figure 1A, Figure 10A to C), this putative ligand binding feature appears ancient in nature, perhaps shared with other PDZ domains that fall into the 5 of 16 defined ligand specificity classes that select for W^-1^ residues [41]. Furthermore, although distinct from erbin in lacking a histidine residue in α1 helix that hydrogen bonds with the ligand T^-2^ side chain (Figure 10G), the unique K_920_ residue in the central region of the *T. adhaerens* Mint α1 helix is predicted to form an analogous interaction with T^-2^ of the ETWC_COOH_ peptide (Figure 10H). This modelled interaction is consistent with the reduction in β-galactosidase activity in the directed yeast 2-hybrid experiments upon mutation of K_920_ to glutamine (Figure 10F), reflecting reduced peptide binding affinity perhaps caused by disrupted hydrogen bonding with T^-2^. Instead, the modelled structure of Mint PDZ-1-CTAV_COOH_ suggests a greater distance between the K_924_ side chain and the carbonyl oxygen of A^-1^, precluding hydrogen bonding, and a clash between side chain sulfur atoms from the C^-3^ residue of the peptide and a methionine residue within the β2 strand of PDZ-1 (M_865_), altogether consistent with our inability to detect an interaction between these two protein elements *in vitro*. In sum, our modelling studies indicate that an interaction between *T. adhaerens* Mint PDZ-1 and the ETWC_COOH_ peptide is at least possible, providing some putative molecular determinants. However, more direct analyses, such as molecular dynamics and/or structural determination, will be required to confirm the molecular nature of this interaction and the various other interactions between Mint proteins and Ca_V_2 channel C-termini.

## Discussion

High-thoughput DNA sequencing has allowed us to learn much about the evolutionary origins of genes that give rise to complexity in animals, including those crucial for electrical and synaptic signaling in the nervous system. Lagging however is our understanding of whether such genes exhibit orthologous protein-protein interactions and cellular functions in the earliest animal lineages, or in pre-metazoan organisms, as defined in bilaterians through wet lab research [96]. One recent pioneering study employed ancetral gene reconstruction to explore the interaction between the third PDZ domain of the postsynaptic scaffolding protein Discs Large Homolgue (DLG) and its binding partner CRIPT, finding it emerged prior to animals in early eukaryotes, but lost binding affinity in select ecdysozoan invertebrates suggesting that even deep ancestral interactions can be subject to dynamic evolution [97]. A few other studies have similarly employed biochemical techniques to explore the origins of select protein-protein interactions *in vitro* [98, 99], or the structure of PDZ-ligand complexes from pre-metazoans [100]. Indeed, continued efforts in this regard will be important as we seek to integrate knowledge about the co-existence of genes in different organismal genomes, with functional data comparing how these genes function in non-mainstream animal systems in particular early diverging animals. This, combined with explorations into the electrophysiological and genetic properties of non-bilaterian neurons and synapses, has the potential to answer important questions about nervous system evolution.

In this study, we explored the evolutionary origins of the first identified interaction between a PDZ domain and a calcium channel C-terminus, Mint and Ca_V_2 [46], and conducted phylogenetic analyses to explore the origins and domain structure of several genes that associate with Mint in bilaterians. One caveat that must be considered when exploring the presence and evolution of genes is that sequences can potentially be missed under a given sequence identification strategy, and future sequencing efforts might uncover homologues in organismal lineages previously reported to lack these genes. Bearing this in mind, our current analysis indicates that Mint/X11 proteins are found broadly in animals, present in all examined animal phyla (Figure 1D). Notably, we used BLAST to search for Mint homologues outside of animals and were unable to identify sequences bearing a conserved PTB/PDZ-1/PDZ-2/P1BM module. The closest sequence with BLAST homology to Mint was from the holozoan eukaryote *Sphaeroforma arctica*, bearing two C-terminal PDZ domains but lacking a PTB domain (NCBI accession number XP_014158662.1). Based on these observations, Mint appears to be unique to animals having evolved via fusion of pre-existing protein domains, a rarer form of gene evolution refered to as type III novelty [101]. Homologues for CASK and Veli, which form a tripartite complex with Mint in vertebrates and bilaterian invertebrates [63–65], were also found broadly in animals, most bearing respective CaMK and L27 domains required for complexing with Mint (Figure 2A and B). However, whether these indeed form complexes in non-bilaterian animals is uncertain given the strong sequence divergence of the Mint N-terminus where the CASK Interaction Domain is found in vertebrates and bilaterian invertebrates (*e.g.*, *D. melanogaster* and *C. elegans* [62]; Figure 1E). In our sequence alignments, we failed to identify conserved signatures corresponding to the CID in non-bilaterian Mint proteins, comprised of an amphipathic alpha helix, a proline-rich linker, and a core binding region bearing an essential tryptophan-valine (WV) motif [62].

Interestingly, we found strong conservation of a YENPTY motif in APP homologues from cnidarians to bilaterians (Figure 2C). This motif is bound by the PTB domain of Mint [50] and is required for membrane internalization and subsequent proteolytic processing of APP into amyloid β (Aβ) [102, 103], a product which is subsequently secreted and accumulates in the brain of Alzheimer’s patients in the form of plaques [104]. Placozoan APP homologues on the other hand lack this motif, while ctenophores appear to lack APP homologues altogether. Instead, all examined animals possess presenilin homologues with conserved C-terminal motifs recognized by both PDZ-1 and PDZ-2 of mammalian Mint 1 and 2 proteins [48, 54, 55] (Figure 2D). This particular interaction links Mint to the production of Aβ, wherein Mint promotes internalization of presenilin (the core subunit of γ-secretase), as well as activity-dependent colocalization of presenilin and APP which presumably precedes cleavage of the latter into Aβ [56, 105]. With respect to neurexin, we identified homologues in most examined animals bearing conserved C-terminal sequences that associate with the Mint-1 PDZ-1 domain from mammals *in vitro* [45, 47] (Figure 3). Mint has been implicated in the proper pre-synaptic localization of neurexin [106], and through its association with presenilin, perhaps also regulates the proteolytic processing of neurexins by this enzyme at synapses [107], two important processes for synaptic homeostasis and function. Indeed, future experiments, for example proteomic analysis of Mint interacting proteins, will be needed to determine whether the various conserved motifs observed in this study indeed mediate interactions with Mint in non-bilaterians and other invertebrates.

This work provides the first experimental evidence that Mint proteins from non-bilaterians can bind Ca_V_2 channel C-termini *in vitro*. First, we identified a robust interaction beween the *N. vectensis* Mint PDZ-1/PDZ-2/P1BM and the Ca_V_2c channel C-terminus, but not the other two isotypes, Ca_V_2a and Ca_V_2b (Figures 6B, G and Figure 8). A major difference in the C-terminal ligands of these channels is that Ca_V_2c bears a threonine at position -2 (*i.e.*, E**T**WC_COOH_), while Ca_V_2 and Ca_V_2b have more canonical Ca_V_2 channel motifs bearing -2 residues of aspartate and glutamate respectively (*i.e.*, D**D**WC and S**E**WC; Figure 1A). In this study, we did not delineate whether the interaction between *N. vectensis* Ca_V_2c and Mint was mediated by PDZ-1 or PDZ-2, since we utilized the full PDZ-1/PDZ-2/P1BM module for all experiments. Our attempts to determine the ligand specificity of PDZ-1 by itself for *N. vectensis* Mint, which would have provided important insights, were unfortunately unsuccesfull. We did however succeed in rendering a bacterial 2-hybrid specificity profile for the *N. vectensis* PDZ-1/PDZ-2/P1BM module, finding a prefence for ligands with tryptophan and valine at positions -1 and 0 respectively (Figure 10C). Hence, the preference for tryptophan at position -1 is consitent with our ability to detect an interaction with the Ca_V_2c ETWC_COOH_ ligand, where perhaps, the threonine at position -2 is permissive for binding while the asparate/glutamate residues in Ca_V_2a and Ca_V_2b disrupt binding. This same module bound the *T. adhaerens* Ca_V_2 ligand sequence of CTAV_COOH_, albeit with lower affinity than ETWC_COOH_ as approximated by β-galactosidase activity (Figure 9), perhaps attributable to the threonine at position -2 and a prefered valine at position 0.

In addition to the *in vitro* interaction between *N. vectensis* Mint and Ca_V_2c, our fluorescence *in situ* hybridization experiments revealed overlapping mRNA expression of these two genes in neurons expressing the marker genes FoxL2 or OtxC [86] (Figure 7). Furthermore, examination of the available single cell RNASeq data for this species confirmed co-expression of these genes in neurons (Figure 8). Thus, it is at least possible that these proteins also physically interact *in vivo*, but future experiments will be required to determine whether this is indeed the case. Expression of Ca_V_2c in *N. vectensis* neurons is consistent with the key role that Ca_V_2 channels play in bilaterian synapses. However, a homologous role for Ca_V_2 channels in presynaptic exocytosis has not been established for cnidarians or ctenophores [36, 108], although fast chemical synaptic transmission in cnidarians requires Ca^2+^ influx [109, 110]. Indeed, while there is general agreement that cnidarian and bilaterian synapses are homologous, ctenophores are proposed to have evolved them independently [60]. Alternatively, synapses evolved once at the base of animals and were lost in placozoans and poriferans (sponges) [111]. For considering these alternate hypotheses, resolving the molecular mechanisms for synaptic trasnmission in cnidarians and ctenophores, including roles for Ca_V_2 channels, will be important. Like Ca_V_2 channels, it is unknown whether Mint plays homologous roles in non-bilaterian neurons as it does in bilaterians, where it is a major player in presynaptic function [57, 112] in part through its selective trafficking of axonal and synaptic proteins [105, 113, 114]. In mice, knockout of the two neuronal Mint genes, Mint-1 and Mint-2, causes lethality in 80% of newborns while the surviving 20% of animals exhibit severe pre-synaptic defects [57]. Neuronal silencing of the two Mint isotypes in *D. melanogaster*, X11α and X11β, also causes larval lethality [113], but not so in *C. elegans* [114], indicating that while Mint is essential in some animals, it appears to play less essential roles in others.

Prompted by our yeast 2-hybrid screens showing that *T. adhaerens* Mint PDZ-1/PDZ-2/P1BM binds several proteins with C-terminal signatures similar to the *T. adhaerens* Ca_V_2 channel sequence of SKCTAV_COOH_ (Figure 1A and Table 1), we conducted directed yeast 2-hybrid experiments revealing that the tandem PDZ module could directly bind the Ca_V_2 C-terminal ligand of CTAV_COOH_ *in vitro* (Figure 9). This finding was consistent with our bacterial 2-hybrid experiments revealing a strong preference for threonine and valine at positions -2 and 0 (Figure 10B). Interestingly, an interaction was not observed for PDZ-1 on its own, making this interaction different from mammals where PDZ-1 mediates binding to Ca_V_2 channels [46]. However, PDZ-1 from *T. adhaerens* Mint did bind the C-terminus of *N. vectensis* Ca_V_2c bearing the ligand sequence ETWC_COOH_ (Figure 9). Furthermore, bacterial 2-hybrid screeing of this domain revealed a preference for tryptophan at position -1 similar to rat Mint-1 PDZ1 (Figure 10A and B), both unexpected findings given that the sequence divergence between *T. adhaerens* and mammalian Mint PDZ-1 falls well below the cut-off that correlates similar ligand specificities between domains [41] (Figures 4 and S1). Altogether, our findings indicate that the placozoan Ca_V_2 channel and Mint can interact *in vitro*, though not through PDZ-1, but rather involving the entire PDZ-1/PDZ-2/P1BM module (Figure 9). This suggests that this non-canonical interaction is being mediated by PDZ-2, or perhaps by a transient locus formed by the association between P1BM and PDZ-1. Consistent with this latter notion is the observation that mutation of K920 to glutamine in PDZ-1, which should have destabilized the PDZ-1-P1BM interaction but minimally impacted PDZ-2 functionality, completely disrupted the PDZ-1/PDZ-2/P1BM interaction with CTAV_COOH_, while maintaining PDZ-1 functionality as demonstrated by its retained ability to bind the *N. vectensis* ETWC_COOH_ ligand (Figure 10). Also, the capacity of Mint to interact with CTAV_COOH_ peptides was also apparent for the cnidarian and ctenophore orthologues, suggesting this mode of binding is conserved. A difference however is that both the cnidarian and ctenophore Mint PDZ-1/PDZ-2/P1BM modules could also bind ETWC_COOH_ peptides, which was not the case for *T. adhaerns* Mint. Although we did not determine the locus for binding of ETWC_COOH_ to cnidarians or ctenophore Mint, our bacterial 2-hybrid data for *N. vectensis* indicates that PDZ-1/PDZ-2/P1BM has a preference for tryptophan at position -1, consistent with weak auto-inihition of PDZ-1 by P1BM.

As noted, placozoans are the most early-diverging animals to possess all three types of metazoan Ca_V_ channel types, Ca_V_1 to Ca_V_3. Functional characterization of these channels revealed general similarities with mammalian and bilaterian orthologues, including the divergent calmodulin feedback regulation that distinguishes Ca_V_1 and Ca_V_2 channels from each other [20, 30, 93]. Unknown however is whether these channels serve similar cellular functions as they do in bilaterians, where for example Ca_V_1 channels are the primary drivers of muscle contraction and Ca_V_2 channels of neurotransmitter release at fast synapses [36, 108]. Furthermore, like cnidarians and bilaterians, the physiological significance of the placozoan Ca_V_2-Mint interaction characterized in this study is unknown. Of note, a recent single cell transcriptome study of four placozoan species, *T. adhaerens*, the closely-related species *Trichoplax* species H2, and two more distantly-related species *Hoilungia hongkongensis* and *Cladtertia collaboinventa*, revealed conserved expression patterns of Ca_V_ channels and Mint, with all three Ca_V_ channel types and the single accessory Ca_V_β subunit of Ca_V_1 and Ca_V_2 channels co-expressed in “peptidergic cells”, a cell type proposed to resemble an evolutionary precursor of neurons and neuroendcorine cells [115]. In the two placozoan species for which Ca_V_2 expression data is available, *Trichoplax* sp. 2 and *C. collaboinventa*, Ca_V_2 expression is mostly restricted to peptidergic cells, mirroring the enriched expression of Ca_V_2 channels in the human brain [116] and Ca_V_2c in *N. vectensis* neurons (Figures 7 and 8). Instead, both the single cell transcriptome data and immunolabeling with polyclonal antibodies against *T. adhaerens* Ca_V_1 [20] revealed broader expression including dorsal epithelial cells which are dynamically contractile and require Ca^2+^ influx to do so [117]. Although the homology of these cells to muscle is unknown, it is notable that broader expression of Ca_V_1 channels oustide of neurons, including muscle, is similarly observed in humans [116]. Lastly, the single cell transcriptome data indicate that Mint and Ca_V_2 are co-expressed in peptidergic cells [115], indicating that the *in vitro* interaction identified here at least have the opportunity to interact *in vivo*, which by extension, provides a putative functional and physiological link between these two proteins.

Our most unexpected finding was that while none of the examined Mint proteins could bind the ctenophore (*M. leydyi*) Ca_V_2 channel C-terminus of SGDI_COOH_ (Figures 1A and 9), *H. californiensis* Mint (PDZ-1/PDZ-2/P1BM) could bind both the *N. vectensis* ETWC_COOH_ and *T. adhaerenes* CTAV_COOH_ peptide ligands with high affinity (Figure 9). This is especially notable given the extreme sequence divergence of both PDZ-1 and PDZ-2 of this species (Figure 4C and D, Figure S1). Indeed, the combined ability of *N. vectensis* and *T. adhaerens* PDZ-1/PDZ-2/P1BM to also bind the non-canonical *T. adhaerens* Ca_V_2 peptide of CTAV_COOH_ suggests this is a conserved and homologous interaction. Altogether, our various observations permit the following three hypotheses about how the interaction between Mint and Ca_V_2 channels evolved (Figure 11): 1) Ca_V_2 channels in cnidarians/bilaterians and placozoans independently evolved the capacity to interact with Mint via emergence of distinct PDZ ligand motifs. This would be consistent with a large-scale analysis indicating that roughly one third of PDZ-ligand pairs observed in humans evolved through emergent C-terminal mutations creating nascent ligands recognized by pre-existing PDZ domains [118]. 2) The placozoan/cnidarian/bilaterian ancestor had a classical type I PDZ ligand of CTAV_COOH_ (*i.e.*, X–S/T-X-Φ_COOH_; where Φ denotes a hydrophobe) [119], and subsequent sequence changes occurred in cnidarians/bilaterians produce rare, non-canonical, DDWC-like ligands (*i.e.*, outside of the 16 specificity classes defined by large-scale analysis [41]). Or 3) the lowest common ancestor of placozoans/cnidarians/bilaterians had a DDWC-like ligand, and placozoans uniquely evolved a type I CTAV_COOH_ ligand.

**Figure 11.**
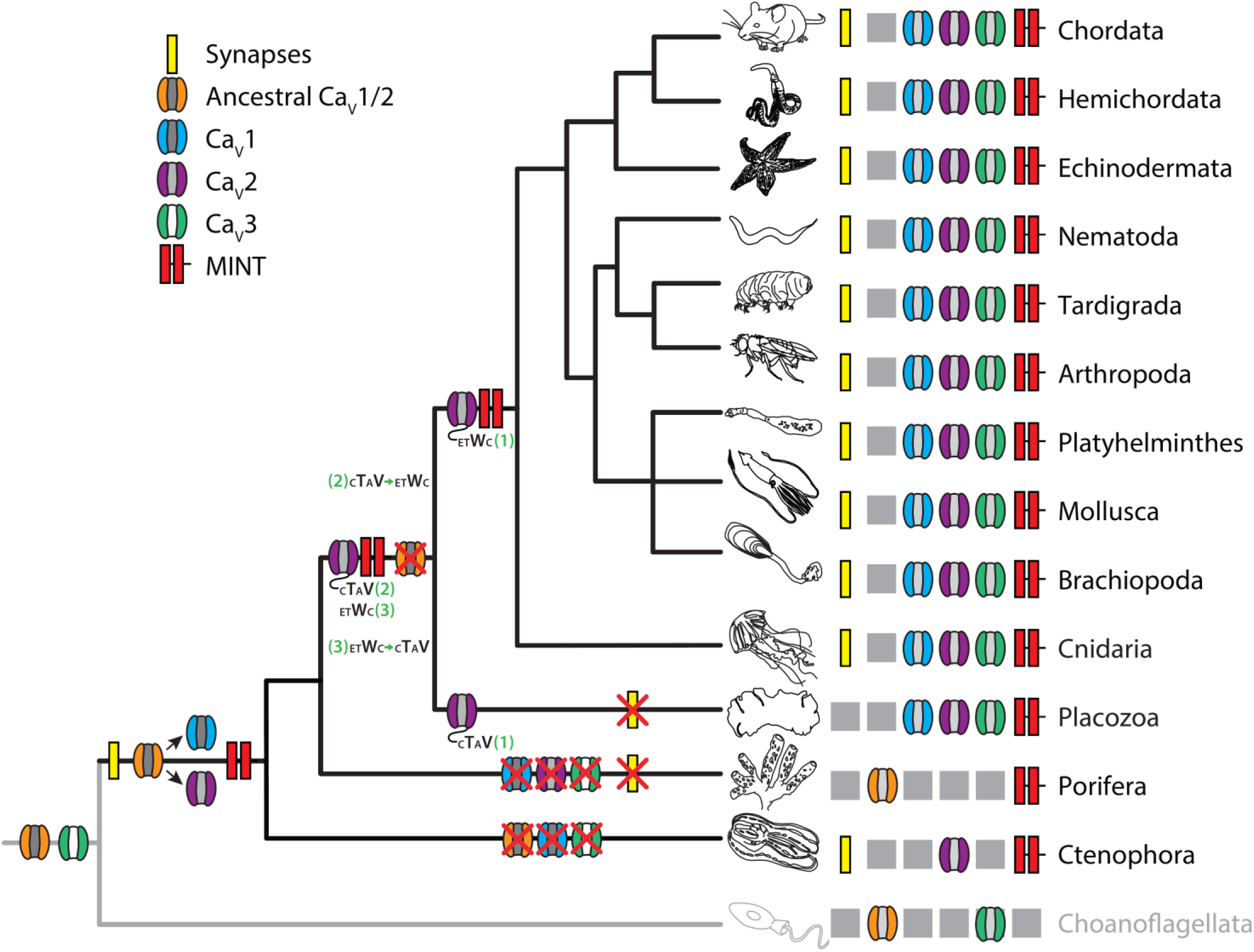
Putative phylogeny of Mint and Ca_V_2 channels in metazoans and choanoflagellates. Animal phylogeny under the ctenophore first and single origin of the nervous system hypotheses, illustrating the known presence/absence of Mint and Ca_V_ channels in different animal phyla and choanoflagellates. The proposed model for Ca_V_ channel evolution is based on previous phylogenetic studies [2–4]. Putative evolutionary events including gains/losses of genes and synapses are represented by symbols on corresponding branches of the tree, while the summary of presence/absence of genes is illustrated bedside the name of each organismal lineage. Under this scenario, Ca_V_1 and Ca_V_2 channels evolved via gene duplication of a Ca_V_1/2 type channel near the stem metazoan lineage [2], along with Mint. This was followed by losses of Ca_V_1/2 channel genes is all animals except poriferans, losses of Ca_V_1 to Ca_V_3 channels in poriferans, and losses of Ca_V_1 and Ca_V_3 channels in ctenophores. The green numbers are used to illustrate three alternative hypotheses for evolution of the Mint-Ca_V_2 interaction as described in the text.

Lastly, it is important consider possible evolutionary interactions between Mint and RIM, since both are known to bind Ca_V_2 channel DDWC_COOH_ and related ligands. As noted, while the physiological relevance of the Ca_V_2-Mint interaction is unknown, RIM plays an essential and conserved role in tethering Ca_V_2 channels at the presynaptic active zone in vertebrates and invertebrates [16, 42–44]. Therefore, a question that emerges is whether the selective pressure to retain DDWC-like SLiMs in the otherwise highly divergent C-termini of cnidarian/bilaterian Ca_V_2 channels came primarily from interactions with RIM, or did the Ca_V_2-Mint interaction also contribute? Without understanding whether the Mint-Ca_V_2 interaction serves some functional purpose *in vivo*, it is not possible to address this question. However, it is notable that in placozoans, an interaction between RIM and Ca_V_2 channels is not expected, since the Ca_V_2 channels have distinct C-terminal motifs that are inconsistent with the known specificity of the RIM PDZ domain. Moreover, the placozoan RIM homologues lack a PDZ domain. Placozoans do possess a second RIM gene which possesses a PDZ domain, referred to as type II RIM [4]. However, these genes are phylogenetically distinct from canonical type I RIMs that interact with Ca_V_2 channels in synapses, and bear highly divergent PDZ sequences making it highly unlikely that they share common ligand preferences with type I RIMs [4]. On the other hand, our study revealed that placozoan Ca_V_2 channels and Mint have the capacity to interact *in vitro*, extending this capability deep into the animal phylogeny, which at least implies that this could be an ancestral interaction with physiological purpose. Lastly, it is noteworthy that none of the examined Mint proteins could bind the ctenophore Ca_V_2 channel C-terminus *in vitro*. Thus, based on our assays, ctenophores seem to distinguish themselves from other animals by lacking the capacity for Mint and Ca_V_2 to interact. Ctenophores also distinguish themselves with respect to RIM, being the only animals with synapses that lack canonical type I RIM, only possessing the divergent type II RIM [4].

## Materials and Methods

### Phylogenetic analyses

Protein sequences used in this study were identified via protein BLAST searches of the NBCI database with Ca_V_2, Mint, CASK, Veli, APP, presenilin, and neurexin orthologues from humans, *C. elegans*, and *D. melanogaster* as queries. The exception are proteins sequences from *T. adhaerens*, *M. leidyi*, and *H. californiensis*, which were obtained from mRNA transcriptomes assembled in house [120], and *H. hongkongensis*, obtained from a published transcriptome [121]. All candidate sequences which produced a BLAST expect value less than 1E-6 were validated as true homologues via reciprocal BLAST of the NCBI database, as well as SmartBLAST [122]. In addition, InterProScan [61] was used to predict and confirm the presence of domains and transmembrane helices expected for the various orthologous proteins (*e.g.*, PDZ domains, SH3 domains, etc.). All extracted and validated sequences used for the presented phylogenetic analyses are provided in separate FASTA files as Supplementary Files 1 (Ca_V_2), 2 (Mint), 3 (CASK), 4 (Veli), 5 (APP), 6 (presenilin) and 7 (neurexin). For consistency, we sought to identify sequences from a finite set of representative species with high quality gene datasets (*i.e.*, possessing full length protein sequences for all/most extracted genes), spanning some of the major animal groupings. In cases where an identified sequence from a given species was too fragmented and missing key domains, an orthologue from a closely-related species was extracted using the same strategy detailed above.

For phylogenetic inference, homologous protein sequences were first aligned with MUSCLE [123], then trimmed with trimAl [124] using gap thresholds ranging from 0.6 to 0.95. The program IQ-TREE 2 [125] was then used to predict Bayesian Information Criterion (BIC) best-fit models and infer maximum-likelihood phylogenetic trees, with 1,000 ultrafast bootstrap replicates to generate node support values. The various phylogenetic trees inferred from different trimmed alignments were then analyzed to select a single tree for each protein type having the highest node support values. All parameters used for phylogenetic inference, including the selected gap thresholds (GT), are as follows: Ca_V_2: GT=0.90, Q.pfam+R7; Mint: GT=0.90, Q.insect+F+R6; CASK: GT=0.70, Q.insect+R4; Veli: GT=0.90, Q.insect+I+G4; APP: GT=0.80, LG+I+R5; presenilin: GT=0.60, Q.pfam+R5; and neurexin: GT=0.90, WAG+F+I+R5 . All aligned and trimmed protein sequences are provided in FASTA format within Supplementary Files 8 (Ca_V_2), 9 (Mint), 10 (CASK), 11 (Veli), 12 (APP), 13 (presenilin) and 14 (neurexin).

### Protein alignments and structural modeling

All protein alignments were prepared using MUSCLE and visualized with Jalview version 2 [126]. For the alignment similarity plots of Ca_V_2 channels and Mint (Figure 1 A and C, respectively), single homologous/paralogous sequences from representative species from each phylum were selected and aligned with MUSCLE, then analyzed with EMBOSS Plotcon [127] using a running windows of 11 amino acids and the EBLOSUM62 comparison matrix. The aligned sequences are provided in FASTA format within Supplementary Files 15 (Ca_V_2) and 16 (Mint). Predictions of MINT protein structure for orthologues from *N. Vectensis* (NCBI accession number XP_032233970.1, amino acids 1095 to 1306), *T. adhaerens* (evg1036658, amino acids 823 to 1034; Supplementary File 2), *H. californiensis* (evg127856, amino acids 337 to 547; Supplementary File 2), *R. norvegicus* (Mint-1, NCBI accession number NP_113967.2, amino acids 630 to 841) comprised of the tandem PDZ-1 and PDZ-2 domains and the P1BM element were generated with Alphafold2 [82] on Compute Canada [128]. The relaxed structural predictions for each homologue bearing the highest per-residue confidence score (pLDDT) were selected and visualized using UCSF ChimeraX [129]. Modelling of the *T. adhaerens* Mint PDZ-1 bound to the ETWC_COOH_ and CTAV_COOH_ sequences was achieved by predicting the structure of the PDZ-1/PDZ-2/P1BM sequence with Phyre2 [130] in the intensive mode (100% of residues modelled at >90% confidence), positioning the peptide based on a peptide-bound structure of the Erbin PDZ domain (PDB ID: 6Q), followed by alteration of the four C-terminal amino acids of the auto-inhibitory sequence (*i.e.*, PEYI_COOH_) to that of the two peptides using Coot [131], then energy minimization with the Chiron Rapid Energy Minimization Server [132]. The resulting structures were then analyzed and visualized with UCSF ChimeraX.

### Fluorescence in situ hybridization

About one hundred seven-day old *N. vectensis* polyps at the tentacle bud stage were transferred into a tube with 1:2 diluted artificial sea water (*Nematostella* medium). To anesthetize/immobilize the polyps, 6.5% MgCl_2_ w/v was added to the water with animals. After one minute spinning at 500 RCF, *Nematostella* medium was replaced with Ca and Mg free seawater (CMFSW) with the following components: NaCl (449 mM), KCl (9 mM), Na_2_SO_4_ (33 mM), and NaHCO_3_ (2.5 mM). Polyps were washed in CMFSW three more times with spinning in between each wash. On the final wash, the polyps were macerated by vigorous pipetting until no big tissue chunks were visible. The dissociated cells were then centrifuged for 5 minutes at 500 RCF and CMFSW was replaced back with *Nematostella* medium. The pellet was then resuspended in *Nematostella* medium and transferred onto cover slips where cells were left to adhere to the cover slips for 1-2 hours. *Nematostella* medium was then removed from the cover slips and these were quickly submerged in 4% PFA in phosphate buffered saline (PBS) for one hour. Fixed cells adhered to cover slips were then rinsed in PBS and dehydrated through an ascending methanol concentration series (30%, 60%, and 100% v/v). Samples were then stored at -20°C in 100% methanol until labelling.

Before labelling, the samples were transferred to ethanol, rinsed one more time with ethanol, and rehydrated in a descending series of ethanol concentrations (100%, 70%, and 50% v/v) in PBS. *In situ* hybridization was then performed with probes and reagents from Advanced Cell Diagnostics, Inc. (Hayward, USA) according to the manufacturer’s instruction. The cells were then pretreated with Protease III diluted 1:15 in PBS before probe application. Sequences of *N. vectensis* gene orthologues were retrieved from NvERTx database [85] and used to generate Z-probes (Nve-Cav2C, #852861; Nve-MINT, #852881-C3; Nve-Otx-C2, #1006821-C2; and Nve-FoxL2-O1-C2, #1007261), which were designed by ACD using the proprietary RNAscope Probe Design pipeline. Special care was taken during the design of *N. vectensis* Ca_V_2c so that the probes do not hybridize with Ca_V_2a and Ca_V_2b transcripts. The negative control included application of 3-Plex Negative Control Probe (#320871), which gave no labelling.

Samples were counterstained with DAPI, mounted in ProLong™ Gold antifade reagent (# P36934, Invitrogen, USA), and examined in LSM 800 confocal microscope with a 63X 1.4 NA objective and fluorescence and DIC optics. Labelling with each probe was done at least twice; the results of independently repeated experiments were similar. For each preparation several randomly chosen fields were scanned. Cell counts were then made for each field, and for each cell in the field we evaluated whether it expressed one gene, combination of any two genes, all three genes, or none of the studied genes. A cell was considered labeled if it possessed at least one fluorescent grain. Mean percent and standard deviation of expressing cells was obtained by averaging the data from all fields. To demonstrate the nature of overlapping expression patterns we drew accurate area-proportional Venn diagrams using eulerAPE software [133]. The fluorescence images were processed with the Fiji image analysis software [134].

### Co-immunoprecipitation experiments

Co-immunoprecipitation experiments were confirmed in triplicate sets of experiments using recombinant proteins transiently expressed in HEK-293T cells (MilliporeSigma, Canada). Briefly, cDNAs corresponding to the C-terminal portions of the *N. vectensis* and *T. adhaerens* Mint PDZ-1/PDZ-2/P1BM protein sequences (as described in the results section) were cloned into the mammalian bicistronic expression vector pIRES2-DsRed2 (Clontech, USA) to express recombinant proteins bearing C-terminal myc epitope tags (Figure 6K). The *N. vectensis* sequence was gene synthesized with codon optimization for mammalian cells, whereas the *T. adhaerens* sequence was PCR amplified from a whole animal cDNA library (all primers and template details for cloning constructs used in these experiments are listed in Supplementary Table 1). Similarly, the entire C-termini of *N. vectensis* Ca_V_2c and *T. adhaerens* Ca_V_2 were PCR amplified from corresponding pIRES2-EGFP plasmid constructs bearing the complete protein coding sequences of the two channels, the former generated via gene synthesis (GenScript), the latter cloned directly from a cDNA library [30], and subsequently restriction digest cloned into the vector pEGFP-C1 (Clontech) to express recombinant fusion proteins bearing an N-terminal Enhanced Green Fluorescent Protein (EGFP) followed by an HA epitope tag (Figure 6J). ΔPDZ ligand variants were created via site directed mutagenesis to introduce premature stop codon upstream of the original stop codon, resulting in the deletion of the 9 most distal amino acids of each Ca_V_2 channel C-terminus. HEK-293T cells were cultured in 25 cm^2^ vented flasks at 37°C in a humidified 5% CO_2_ incubator in 6 mL of DMEM supplemented with 10% FBS and 0.5 mL of penicillin/streptomycin (Sigma-Aldrich, Canada; 5,000 units/mL penicillin and 5 mg/mL streptomycin). For transfections, cells were grown to 60% confluency then transfected with 3 μg of each relevant DNA construct using the PolyJet^TM^ transfection reagent (SignaGen Laboratories, USA). 6 hours later, cells were washed with warm 1x Phosphate Buffered Saline (PBS) (Gibco, ThermoScientific Scientific, Canada) and placed at 37°C for 24 to 36 hours. Cells were then transferred to a 28°C 5% CO_2_ incubator for 2 to 3 days to increase protein expression [135], then harvested by first washing with 1x PBS, then solubilizing in extraction buffer containing 0.5% Triton X-100, 20 mM imidazole at pH 6.8, 100 mM NaCl, 1 mM EDTA, 1 mM DDT, and 1x Halt™ Protease Inhibitor Cocktail (ThermoFisher Scientific). Lysates were centrifuged at 16,000 × g for 15 min and treated with 1 unit/μL Benzonase Nuclease (E1014, Millipore Sigma, Canada) for 30 minutes on ice. Lysates were then then quantified for total protein content using the Pierce^TM^ BCA Protein Assay Kit (ThermoFisher Scientific) then either snap frozen in liquid nitrogen and stored at -80 °C or immediately used in downstream experiments.

Co-immunoprecipitation experiments were performed using the Pierce™ Crosslink IP Kit (ThermoFisher Scientific) with minor modifications to the manufacturer’s protocol. Briefly, the included Pierce™ Protein A/G Plus agarose beads were incubated with rabbit anti-myc polyclonal antibody (ab9106, Abcam, USA) and cross-linked using 2.5 mM DSS crosslinker. Protein lysates were then pre-cleared of non-specificities using the Pierce™ Control Agarose Resin and incubated with the cross-linked Pierce™ Protein A/G Plus Agarose beads. The beads were then washed with extraction buffer multiple times and immunoprecipitated complexes were boiled off in NuPAGE™ 4x Sample Loading Buffer (ThermoFisher Scientific, Canada). The co-immunoprecipitated and immunoprecipitated proteins were then characterized by Western Blotting using anti-HA monoclonal antibody (ab18181, Abcam, USA) and anti-myc monoclonal antibody (ab18185, Abcam, USA), respectively, both at a dilution of 1:1000. Input was 1% of the amount of cell lysate used for co-immunoprecipitation and was similarly characterized by Western blotting with anti-myc and anti-HA monoclonal antibodies. Chemiluminescent signals on blots were detected using SuperSignal West Femto Maximum Sensitivity Substrate (Millipore Sigma) imaged an ImageQuant LAS 500 Chemiluminescence CDD Imager (GE Life Sciences, USA). All raw images of Western blots are provided in Supplementary File 17.

### Yeast 2-hybrid and bacterial 2-hybrid experiments

The *T. adhaerens* yeast 2-hybrid cDNA library was constructed using the Make Your Own “Mate & Plate^TM^” Library System (Takara Bio USA, Mountain View, CA), using whole animal total RNA extracted from approximately 30 animals using a Nucleospin RNA Plus Mini Kit (Macherey-Nagel, Düren, Germany), both according to manufacturer’s instructions. The resulting yeast 2-hybrid library was screened according to the manufacturer’s instructions, using the Mint PDZ-1/PDZ-2, P1BM and PDZ-1 proteins sequences as bait cloned expressed from the bait vector pGBKT7 as fusion proteins N-terminally tagged with the Gal4 DNA binding domain. All primers used for PCR amplification and restriction digest cloning, as well as template details, are provided in Supplementary Table 1. For the directed yeast 2-hybrid screens, the 70 most distal amino acids of *T. adhaerens* Ca_V_2 and *M. leidyi* Ca_V_2 were PCR amplified from previously cloned pIRES2-EGFP plasmid constructs containing the full-length protein coding sequences of each channel, then cloned into the prey vector pGADT7 with and without the last four amino acids of the channel C-termini to express fusion proteins with the Gal4 activation domain at the N-terminus. The C-terminus of *N. vectensis* Ca_V_2c was similarly cloned, using a gene synthesized coding sequence (GenScript, USA) as template for the PCR reaction. All yeast 2-hybrid screens were done using the Matchmaker® Gold Yeast Two-Hybrid System (Takara Bio USA) according to the manufacturer’s instructions, including necessary controls to rule out auto-activation of protein-protein interaction reporter genes by the various Mint insert proteins. Measurements of β-galactosidase reporter gene activity were made with the ThermoScientific Yeast β-Galactosidase Assay Kit according to the manufacturer’s instructions, on flat bottom 96-well plates read using a BioTek Epoch Microplate Spectrophotometer (Agilent Technologies Canada Inc., Canada) to determine cell densities (OD_600_) and colorimetric enzyme activity (absorbance at 420 nm).

### Bacterial 2-hybrid experiments

Bacterial 2-hybrid assays were carried out as previously described [94] with slight modifications to the assay. First, we modified the peptide length of the library from 7 amino acids to 6, finding that in general screens of PDZ-peptide specificity rarely enrich any preference at position -6. Reducing the peptide length in this way reduced the complexity of the library, which permitted us to sample potential ligand sequences in greater depth. We used the same library building approach as previously described to minimize bias in the library [136]. Second, we carried out all selections on agar plates containing 5 mM 3-aminotriazole (3AT), an inhibitor of our HIS3 reporter which minimizes the enrichment of false positives. We used plates rather than liquid culture as we found similar data was produced with significantly less effort. Once grown, the surviving members of the library were pooled, sequenced by Illumina, and motifs generated as previously described [94].

### Availability of Data and Materials

All datasets of protein sequences used for phylogenetic and/or in silico structural analyses are included in the published article and its supplementary information files. All other datasets used and/or analysed during the current study available from the corresponding author on reasonable request.

## Acknowledgements

We would like to thank the Light Imaging Facility and Dr. Carolyn Smith of the National Institute of Neurological Disorders and Stroke, Drs. Kristen Koenig and Kyle McCulloch for providing the *N. vectensis* specimens used in this study, Dr. Elise F. Stanley for her feedback during early stages of this study, and Dr. Joseph Ryan for his feedback on the manuscript.

## Funding

This research was funded by an NSERC Discovery Grant (RGPIN-2021-03557), an NSERC Discovery Accelerator Supplement (RGPAS-2021-00002), an Ontario Early Researcher Award (ER17-13-247), and a Canadian Foundation for Innovation Grant (35297) to A. Senatore, Ontario Graduate Scholarships to A. Singh, W. E., and G. M., an NSERC Canadian Graduate Scholarship to J. G., an NSERC Discovery Grant (RGPIN-2023-05615) and a Canadian Foundation for Innovation Grant (40684) to M. A. C., and a NIH grant (R01GM133936) to M. N..

## Notes

### Competing Interest Statement

The authors have declared no competing interest.

### Summary of Updates

1. New phylogenetic analyses were done to incorporate alternate node support values and several additional neurexin sequences from non-bilaterians missed in our previous analysis. 2. N. vectensis single cell RNA-Seq data was mined to corroborate the in situ hybridization experiments.

